# An inappropriate decline in ribosome levels drives a diverse set of neurodevelopmental disorders

**DOI:** 10.1101/2024.01.09.574708

**Authors:** Chunyang Ni, Leqian Yu, Barbara Vona, Dayea Park, Yulei Wei, Daniel A Schmitz, Yudong Wei, Yi Ding, Masahiro Sakurai, Emily Ballard, Yan Liu, Ashwani Kumar, Chao Xing, Hyung-Goo Kim, Cumhur Ekmekci, Ehsan Ghayoor Karimiani, Shima Imannezhad, Fatemeh Eghbal, Reza Shervin Badv, Eva Maria Christina Schwaibold, Mohammadreza Dehghani, Mohammad Yahya Vahidi Mehrjardi, Zahra Metanat, Hosein Eslamiyeh, Ebtissal Khouj, Saleh Mohammed Nasser Alhajj, Aziza Chedrawi, César Augusto Pinheiro Ferreira Alves, Henry Houlden, Michael Kruer, Fowzan S. Alkuraya, Can Cenik, Reza Maroofian, Jun Wu, Michael Buszczak

**Author notes:** Equal contribution (Chunyang Ni, Leqian Yu, Barbara Vona).

## Abstract

Many neurodevelopmental defects are linked to perturbations in genes involved in housekeeping functions, such as those encoding ribosome biogenesis factors. However, how reductions in ribosome biogenesis can result in tissue and developmental specific defects remains a mystery. Here we describe new allelic variants in the ribosome biogenesis factor *AIRIM* primarily associated with neurodevelopmental disorders. Using human cerebral organoids in combination with proteomic analysis, single-cell transcriptome analysis across multiple developmental stages, and single organoid translatome analysis, we identify a previously unappreciated mechanism linking changes in ribosome levels and the timing of cell fate specification during early brain development. We find ribosome levels decrease during neuroepithelial differentiation, making differentiating cells particularly vulnerable to perturbations in ribosome biogenesis during this time. Reduced ribosome availability more profoundly impacts the translation of specific transcripts, disrupting both survival and cell fate commitment of transitioning neuroepithelia. Enhancing mTOR activity by both genetic and pharmacologic approaches ameliorates the growth and developmental defects associated with intellectual disability linked variants, identifying potential treatment options for specific brain ribosomopathies. This work reveals the cellular and molecular origins of protein synthesis defect-related disorders of human brain development.

**Highlights:**

- *AIRIM* variants reduce ribosome levels specifically in neural progenitor cells.
- Inappropriately low ribosome levels cause a transient delay in radial glia fate commitment.
- Reduced ribosome levels impair translation of a selected subset of mRNAs.
- Genetic and pharmacologic activation of mTORC1 suppresses AIRIM-linked phenotypes.

## Introduction

Human brain development depends on the coordinated action of various signaling pathways, coupled with cell-specific and stage-specific gene expression, to specify a diverse range of cell types ^1^. During the initial stages of brain development, neuroepithelial cells undergo proliferation and differentiate into radial glial progenitors, intermediate progenitors, and outer radial glial progenitors. Progenitor cells in the dorsal region differentiate into excitatory neurons, while those in the ventral region give rise to interneurons that subsequently migrate to the dorsal cortex. It is well known that initial cell fate decisions are influenced by the differential expression of crucial transcription factors ^2,3^. However, the role of post-transcriptional regulation in early brain development remains less explored.

The study of neurodevelopmental disorders (NDDs) has provided key insights into the mechanisms that govern these early steps in human brain development. NDDs are a broad spectrum of childhood disorders that affect more than 4.7% of children globally, and include intellectual disabilities, seizures, defects in sensory perception, and in extreme cases, microcephaly ^4^. NDDs are most frequently caused by genetic lesions. However, only a small proportion of these mutations disrupt genes that have a primary role in human brain development ^5^. Rather, a significant number of NDD-associated allelic variants map to genes involved with general cellular housekeeping activity. How mutations in housekeeping genes can result in both tissue and developmentally specific phenotypes is unclear.

Mutations in ribosomal protein and ribosome biogenesis genes cause a group of related diseases called ribosomopathies, which include Diamond Blackfan Anemia (DBA), Treacher Collins Syndrome, X-linked dyskeratosis congenita (DC), and cartilage hair hypoplasia (CHH) ^6,7^. The phenotypes associated with ribosomopathies vary widely. For example, patients with DBA, which is linked with mutations in several genes including RPS19, RPL5, and TSR2, primarily present with severe anemia, and only occasionally suffer from intellectual disabilities ^8^. By contrast, TCS, which is associated with loss of the Pol I factor TCOF1, results in craniofacial malformations ^9^. Recent results indicate that mutations in Ribosomal RNA Processing 7 Homolog A, RRP7A and dysfunction of ribosome quality control (RQC) mechanisms are also associated with neurological disorders ^10,11^. Disruption of ribosome biogenesis or function often results in a nucleolar stress response that induces p53 activity. Genetic ablation of p53 can suppress many of the phenotypes associated with models of ribosomopathies ^12–14^. However, the molecular and cellular mechanisms responsible for the tissue specificity of these diseases remains incompletely understood.

Recently, we have characterized a complex, composed of AIRIM (previously named C1orf109), AFG2A (SPATA5), AFG2B (SPATA5L1) and CINP, that promotes the recycling of RSL24D1 from cytoplasmic pre-60S ribosomal subunits back to the nucleolus ^15^. AFG2A is the human ortholog of yeast Drg1, which plays a well characterized role in ribosome biogenesis ^16^. Additional studies have further implicated human AFG2A and AFG2B in ribosome production ^17,18^. Strikingly, human genetic studies have identified a growing number of allelic variants in AFG2A, AFG2B, and CINP that are specifically associated with a range of NDDs, including intellectual disabilities, seizures, hearing loss, and microcephaly ^19–25^. Here, we identify new variants in AIRIM associated with a similar range of neurological phenotypes. Thus, unlike variants linked with DBA and other ribosomopathies, disruptions in the AIRIM complex primarily result in NDDs. These observations beg the question of whether these mutants represent a new class of ribosomopathy, and if so, how do disruptions in 60S biogenesis specifically result in brain specific abnormalities.

Specific NDDs are not always recapitulated in mouse models. To overcome this experimental barrier, many groups have turned to using human iPSC derived brain organoids ^26–28^. These organoid models have proven useful for studying human brain development. Transcriptional and epigenetic analyses indicate that brain organoids in 3D culture recapitulate many of the same developmental processes and gene expression programs that occur during fetal development in vivo ^2,29^. Moreover, these models have been extremely useful for characterizing mechanisms involved in various neurodevelopmental diseases including microcephaly and autism ^28,30^.

Generating complex brain-like organoids from human induced pluripotent stem cells (iPSCs) provides us with an opportunity to study the mechanisms by which mutations in the AIRIM complex affect human brain development. The characterization of these variants using cerebral organoids provides evidence that the dynamic regulation of ribosome levels is critical for early neurodevelopment. Defects associated with patient variants in both AIRIM and AFG2B can be traced back specifically to neuroepithelial differentiation. These variants cause reductions in the translation of a specific subset of mRNAs encoding both components of protein synthesis machinery and factors that direct early cell fate specification. This decrease in the protein expression of specific factors causes a delay in the neuroepithelial to radial glial cell transition. Genetically or pharmacologically increasing mTOR signaling alleviates the growth, enhanced cell death, and cell-fate specification phenotypes observed in organoids that carry a pathogenic *AIRIM* variant. These findings highlight the importance of the stage specific regulation of ribosome availability in the developing human nervous system.

## Results

### Patients with AIRIM variants display central nervous system abnormalities

How mutations in ribosomal proteins and ribosome biogenesis factors disrupt the development and function of some specific tissues but do not impact others remains an open question. We reasoned that identifying new variants in ribosome biogenesis factors specifically linked with NDDs would provide further insights into how perturbations in translation can selectively disrupt early brain development. Pedigrees, clinical details, and variant characteristics identified a cohort comprised of seven families with similar NDDs (**Figure 1A; Figure S1; Tables S1 and S2**). Affected individuals presented with severe-to-profound global developmental delay/intellectual disability (11/11) and never achieved developmental milestones. The majority of individuals from whom information is available concomitantly show muscular hypotonia accompanied with limb spasticity and dystonia (each 8/8), microcephaly (8/10), as well as hearing (6/7) and vision impairment (6/8) and dysmorphism (**Table S1**). Infantile seizures were reported in seven individuals and further characterized in six as generalized tonic and clonic (2/6), myoclonic (2/6), tonic-clonic (1/6) and infantile spasm (1/6). The neuroimaging analysis revealed severe supratentorial brain atrophy with diffusely thin corpus callosum, and ex-vacuum dilatation of the lateral ventricles along with under opercularization of the Sylvian fissures, while the brainstem and cerebellum were relatively preserved (**Figure 1B; Tables S1, S2**). Additionally, abnormal signal intensity was observed in both the deep and superficial white matter, indicating impaired myelination with hypomyelination appearance. However, normal myelination was observed in the limbs of the internal capsules, optic tracts, and posterior fossa structures.

**Fig. 1.**
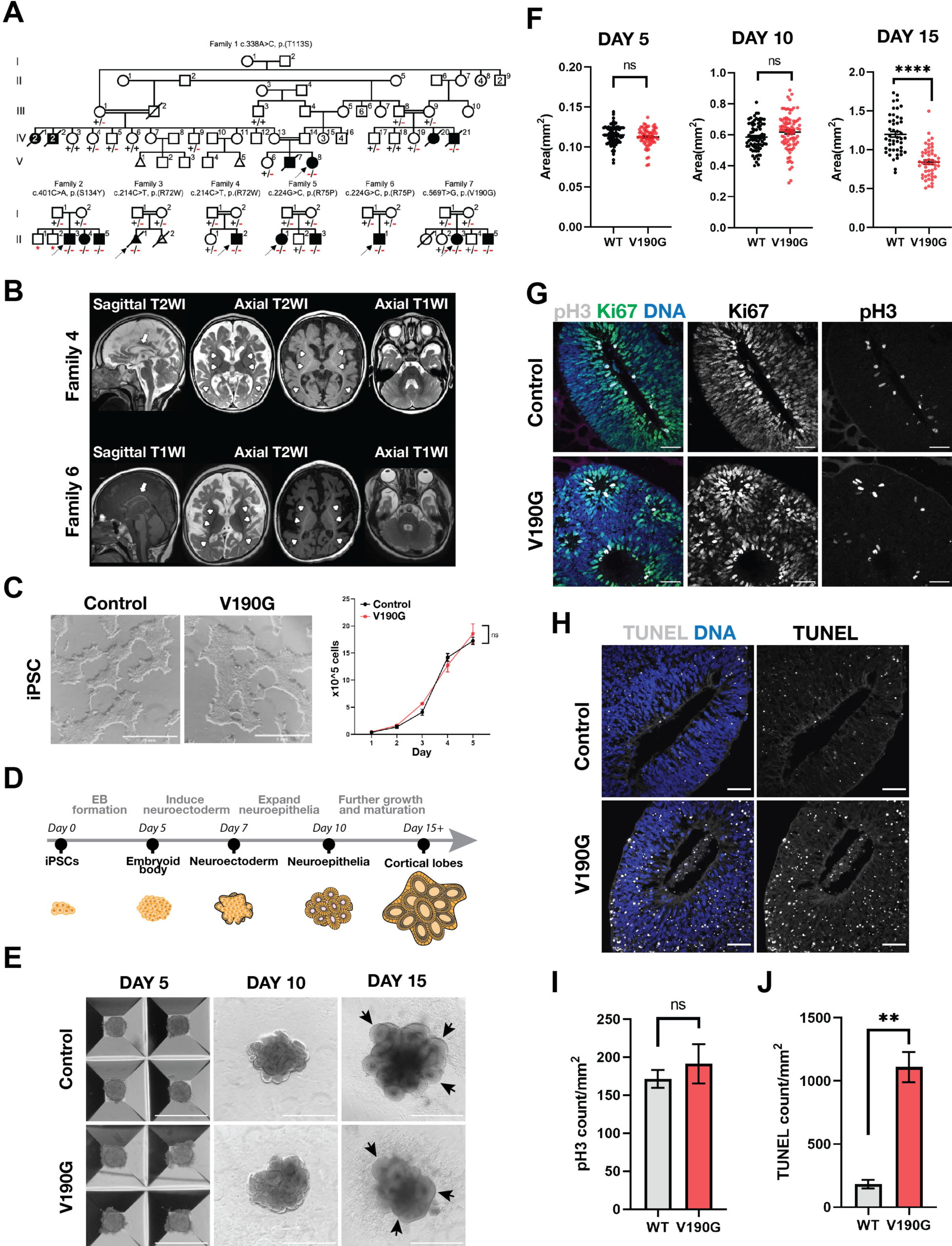
The NDD associated *AIRIM^V^*^190^*^G^* variant causes growth defects at a specific stage of cerebral organoid development. **(A)** Pedigrees showing segregation of the variants. Affected and unaffected individuals are indicated by filled and open and filled symbols, respectively. Probands are marked with arrows. Double lines indicate consanguinity. Red asterisks represent a non-homozygous genotype. One affected individual (Family 3, II:1) was medically aborted at 18 weeks subsequent to the identification of abnormal brain development via basic sonography. An affected cousin in Family 1 (IV:21) passed away following study enrollment with a remarkable family history showing six other similarly affected individuals who had previously deceased before initiation of the study. (**B**) Brain MRI of the proband in family 4 (II:1) at 1 year 11 months sagittal T2-weighted image (WI), axial T2WI and axial T1WI; brain MRI in family 6 (II:1) at 6 months old; sagittal T1WI, axial T2WI and axial T1WI. Severe atrophy in the supratentorial compartment of the brain, including a diffusely thin corpus callosum (arrows) and diffuse enlargement of the extra-axial spaces with bilateral under opercularization of Sylvian fissures. Abnormal T2WI hyperintensity of the deep and superficial white matter with relatively normal myelination appearance along the anterior and posterior limbs of the internal capsules, as well as the optic tracts (arrowheads). It is also important to note the relatively normal appearance of the overall structures of the posterior fossa with normal volume, morphology, and myelination appearance of the cerebellum. **(C)** Brightfield images of control and *AIRIM^V^*^190^*^G^* iPSCs and a direct comparison of their corresponding proliferation under normal culture conditions. (**D**) Schematic of the timeline for generating brain organoids from pluripotent stem cells. (**E**) Bright field images of control and mutant (V190G) organoids at days 5, 10, and 15. Black arrows indicate neuroepithelial buds, which appear less organized and elongated in the mutant at day 15. Scale bar, 1mm. (**F**) Quantification of bright field images at days 5 (left), 10 (middle) and 15 (right) show control neural tissue is enlarged relative to mutant (V190G); ∗∗∗∗p < 0.0001, Unpaired t test with Welch’s correction, n (Day 5) = 73 control EBs 62 mutant EBs from 3 independent batches, n (Day 10) = 77 control organoids 84 mutant organoids from 4 independent batches, n (Day15) = 46 control organoids 51 mutant organoids from 3 independent batches, error bars are SEM. **(G)** Representative images showing the proliferation marker phospho-Histone 3(pH3, grey) and Ki67(green) signal in control (WT) and mutant (V190G) day 15 organoids. Scale bar, 50 μm. (**H**) Representative images showing TUNEL labeling in control and mutant (V190G) day 15 organoids. Note mutant organoids exhibit significantly more TUNEL signal. Scale bar, 50 μm. (**I**) Quantification of pH3 signal of individual neuroepithelial bud at day 15 control (WT) and mutant (V190G) organoids. Unpaired t test with Welch’s correction. n = 12 control and 11 mutant imaged regions from 2 independent batches. error bars are SEM. (**J**) Quantification of TUNEL signal of individual neuroepithelial bud showing mutant organoids exhibit more TUNEL signal. TUNEL counts were normalized to the area of the imaged bud. ∗∗∗∗p < 0.0001, Unpaired t test with Welch’s correction. N = 6 control and 6 mutant imaged regions from 2 independent batches. error bars are SEM.

Using exome sequencing and homozygosity mapping, we identified a homozygous missense variant c.338A>C; p.(Tyr113Ser) in *AIRIM* residing in a ∼12 Mb run of homozygosity. Continued sequencing of either proband or parent-child trios identified four other homozygous missense variants: c.401C>A; p.(Ser134Tyr); c.214C>T; p.(Arg72Trp); c.224G>C; p.(Arg75Pro); c.569T>G; p.(Val190Gly) (**Figure 1A**; **Table S1; Figure S1**). All variants segregated within the families, and they are either totally absent or present in an extremely low allele frequency in sequence variant databases. Recent results indicate that AIRIM functions with AFG2A, AFG2B and CINP to promote the recycling of RSL24D1 from cytoplasmic pre-60S ribosome subunits back to the nucleolus. Of note, variants in these three other complex members are associated with a strikingly similar range of neurodevelopmental defects^19–22,24,25,31^ (**Figure S1**), indicating these phenotypes linked to all four genes likely originate from disruption of a common function.

### Organoids as model of brain ribosomopathies

To investigate how variants in *AIRIM* cause specific defects in human brain development, we introduced the Val190Gly (V190G) mutation into iPSCs, generated cerebral organoids from mutant and isogenic control iPSCs, and tracked organoid growth through five stages: undifferentiated iPSCs (day 0), embryoid bodies (EBs) (day 5), organoids composed mostly of epithelial neural stem cells (day 10); initiation of cortical lobe formation and neurogenesis (day 15), and initiation of cortical plate formation (day 30) (**Figure 1C,D**). In parallel, we generated patient specific iPSCs from fibroblasts transheterozygous for the Ile466Met and Val245Glu mutations in *AFG2B* ^24^ and parental controls. We subjected these iPSCs to the same differentiation protocol. The *AIRIM^V^*^190^*^G^*mutation did not affect proliferation or pluripotency of starting iPSCs (**Figure 1C; Figure S2A**). Similar results were obtained for *AFG2B* patient derived iPSCs (**Figure S2B,C**). During organoid formation, *AIRIM^V^*^190^*^G^* and *AFG2B^I^*^466^*^M/V^*^245^*^E^*mutant iPSCs differentiated into day 5 EBs and day 10 neural epithelia without obvious defects (**Figure 1E,F; Figure S3C**). By contrast, *AIRIM^V^*^190^*^G^*and *AFG2B^I^*^466^*^M/V^*^245^*^E^* organoids exhibited reduced size and less elongated neuroepithelia buds relative to controls on day 15 (**Figure 1E,F; Figure S3C**).

We reasoned that the observed differences in organoid size between controls and allelic variant organoids could arise from increased cell death, decreased cell proliferation, or both. Staining of sectioned day 15 organoids revealed comparable levels of cell proliferation (pH3 and Ki67), but significantly increased cell death (TUNEL) in mutant *versus* control organoids (**Figure 1G-J; Figure S3D-G**). Interestingly, this phenotype did not lead to progressive degeneration, as mutant organoids continued to develop and formed cortical plate on day 30, albeit at a smaller size (**Figure S3A,B)**.

### Ribosome and protein synthesis are dynamically regulated in early cortical development

Previous studies indicated that loss of *AIRIM* or *AFG2B* results in defects in late 60S maturation, marked by disruption of Ribosomal L24 Domain Containing 1 (RSL24D1) recycling off of pre-60S subunits back to the nucleus^15^. Organoids carrying the *AIRIM^V^*^190^*^G^* mutation exhibited a modest but significant decrease in nuclear RSL24D1 (**Figure S4**), suggesting reduced levels of ribosome biogenesis. To characterize how the *AIRIM^V^*^190^*^G^* variant affects global protein levels, we performed two orthogonal assays: O-propargyl puromycin (OPP) labeling and quantitative tandem mass tag mass spectrometry (TMT-MS) analysis (**Figure 2A**). OPP is an analog of the tyrosyl-tRNA mimic puromycin and can be used to pulse label nascent peptides to measure translation activity (**Figure 2A**)^32,33^. We performed OPP labeling on control and mutant organoids on days 5, 10 and 15 of differentiation (**Figure 2B**). Mutant organoids exhibited similar levels of protein synthesis on day 5 when compared to the controls. By contrast, OPP labeling in mutant organoids was significantly reduced relative to control organoids on days 10 and 15 (**Figure 2B,C**).

**Figure 2.**
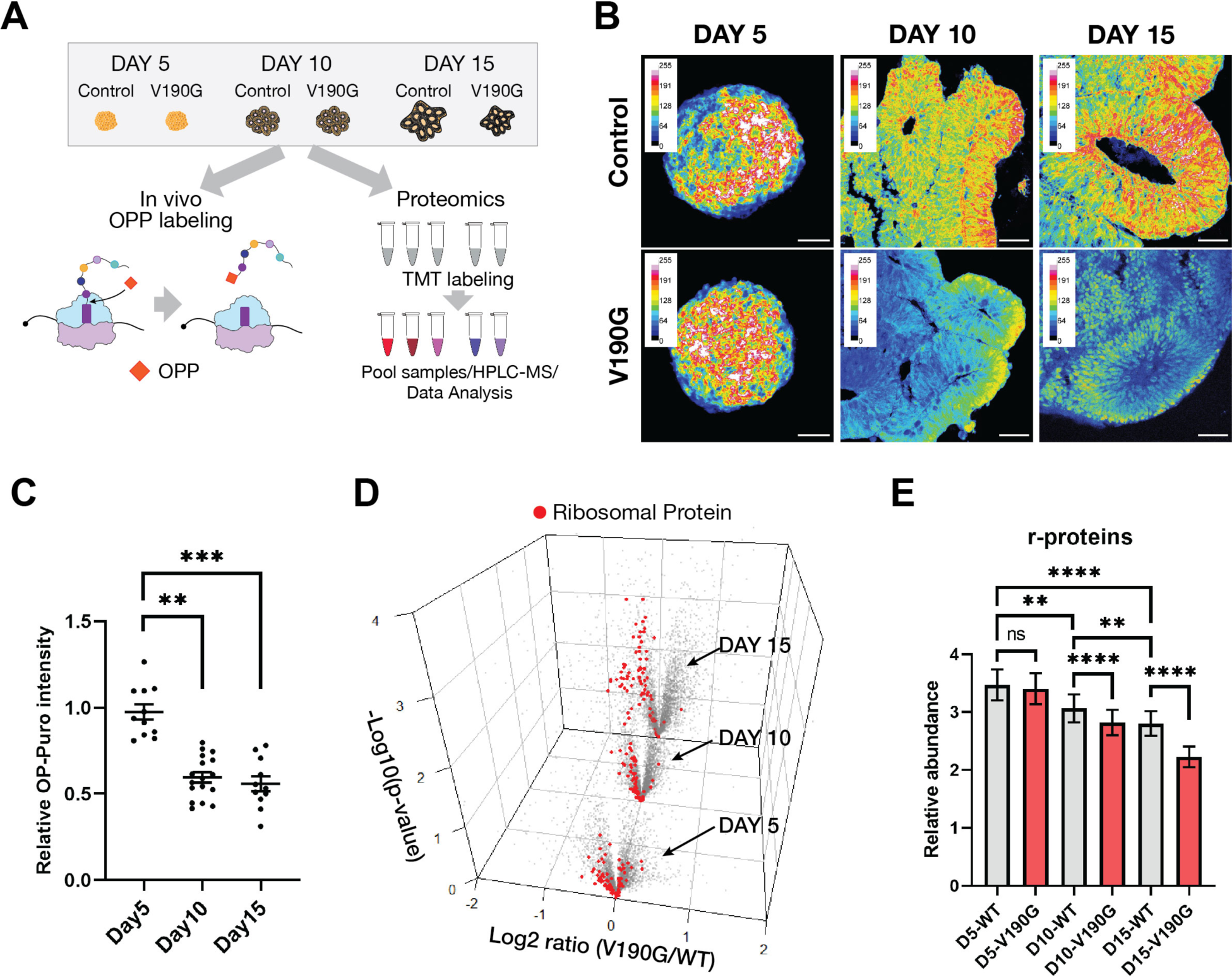
The *AIRIM^V^*^190^*^G^* variant causes stage-specific defects in protein synthesis and ribosome levels. (**A**) Experimental schemes of OP-puro based translation capacity and TMT assays. (**B**) Representative images showing the OP-puro signal in control of control and mutant (V190G) organoids at days 5, 10, and 15. Note mutant organoids exhibit significantly lower OP-puro signal than control starting at day 10. Scale bar, 50 μm. (**C**) Quantification of OPP signal of individual neuroepithelial bud at day 5, 10 and 15 mutant (V190G) organoids relative to their corresponding control. Dunn’s multiple comparisons test. n = 11, 17, 11 individual EB or neuroepithelial buds imaged from 2 independent batches. error bars are SEM. (**D**) Volcano plots of the −log 10 -transformed P value versus the log 2-transformed ratio of Mutant/Control organoids, at days 5, 10, and 15 (n = 3539, 3544, 3541 proteins, n = 3 biologically independent samples). Discovery determined using the Two-stage linear step-up procedure of Benjamini, Krieger and Yekutieli. (**E**) Mean ratio values (± SD) for r-proteins as in h (n = 76 r-proteins; n = 3 biologically independent samples). Adjusted P values were calculated from Tukey’s multiple comparisons test.

We next performed TMT-MS on day 5, 10 and 15 control and *AIRIM^V^*^190^*^G^* organoids to systematically examine the abundance of ribosome components during organoid formation (**Figure 2D,E; Table S3**). During the formation of control organoids, we observed that ribosomal protein levels declined between days 5 and 15, suggesting that changes in ribosome levels are part of the normal differentiation program (**Figure 2D,E**). By comparison, although *AIRIM^V^*^190^*^G^*organoids on day 5 exhibited comparable ribosome protein levels, they showed a more dramatic reduction on days 10 and 15. These findings further suggest that the *AIRIM^V^*^190^*^G^*variant compromises ribosome formation and global protein synthesis, prior to when growth defects first manifest during brain organoid formation.

### Delayed radial glia (RG) differentiation in organoids with lower protein synthesis

To study how decreases in ribosome levels and mRNA translation influence organoid differentiation dynamics, we performed scRNA-seq on control and *AIRIM^V^*^190^*^G^*mutant organoids across five different time points (**Figure 3A,B**). The sequencing data of each genotype and time point were integrated using Harmony and visualized by uniform manifold approximation and projection (UMAP) embedding^34^. These integrated data enabled a comprehensive assessment of transcriptional dynamics during organoid formation. The developmental progression and gene expression programs of our control organoids were consistent with other studies^30,35,36^. As expected, organoid differentiation proceeded from iPSCs (POU5F1) to neural progenitors (PAX6, VIM), then to intermediate progenitors (EOMES) and neurons (DCX, TUBB3, MAP2). We also observed a small neural crest/mesenchymal population (DCN, LUM) as well as a population that resembles cells within the choroid plexus (TTR) (**Figure 3C**).

**Figure 3.**
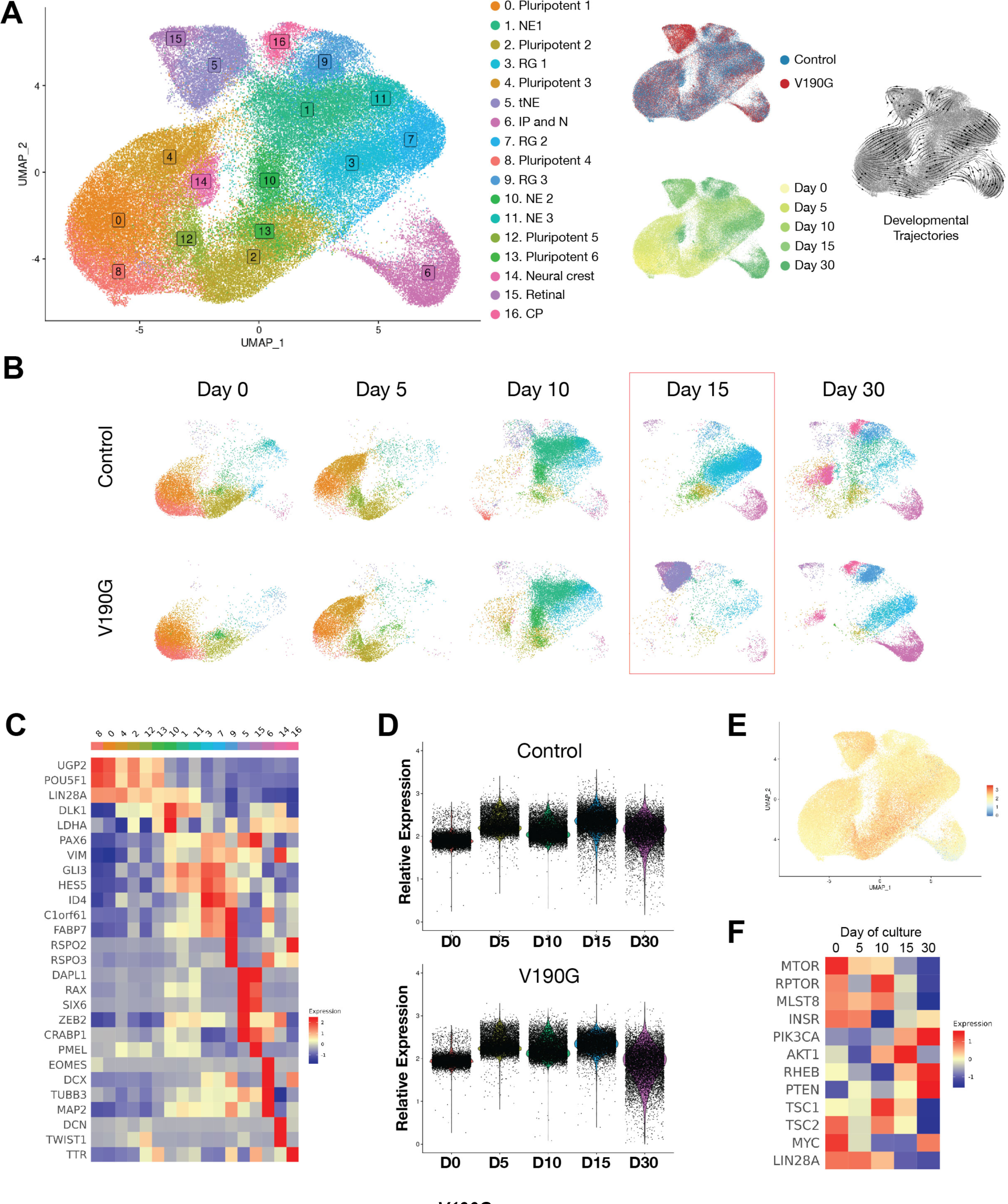
scRNA-seq reveals *AIRIM^V^*^190^*^G^* mutant organoids display a transient delay of neuroepithelial differentiation. (**A**) UMAP embedding of full integrated dataset, colored by Louvain clusters, time point, genotype and RNA velocity. NE, Neuroepithelium; tNE, transitioning NE; RG, radial glia; IP, intermediate progenitor; N, neuron; CP, choroid plexus. (**B**) Split-view of full integrated dataset, colored by Louvain clusters. (**C**) Heatmap showing the average expression of cluster marker genes related to brain development from the full integrated dataset. (**D**) Gene module expression scores for a set of 79 r-protein genes across developmental time-points, in cells from control (WT) samples and from mutant (V190G) samples. **(E)** Gene module expression scores for a set of 79 r-protein genes across developmental time-points. (**F**) Heatmap showing the average expression of translation regulator genes across the development of control organoids.

The single cell transcriptomes of control and *AIRIM^V^*^190^*^G^* iPSCs and day 5 EBs largely overlapped with one another (**Figure 3A,B**). Despite the observed differences in protein synthesis levels (**Figure 2B,C**), the transcriptional states of neuroepithelia (NE, LDHA) and transitioning neuroepithelia (CDH2, VIM, ZEB2) in both control and mutant day 10 organoids were also similar (**Figure 3A-C)**. However, while the neuroepithelia in control organoids continued to differentiate into radial glial progenitors (RGs, FABP7/BLBP), accompanied by signs of neurogenesis on day 15, mutant organoids exhibited delays in RG fate specification and less neurogenesis (**Figure 3B**, red box). Instead, most progenitors in day 15 *AIRIM^V^*^190^*^G^* organoids continued to express ZEB2 (cluster 5), a regulator that functions during the NE to RG transition, indicating that while mutant NE cells initiated RG specification they did not complete this transition in a timely manner. Despite this delay, however, on day 30, both control and mutant organoids contained an abundant number of committed RGs and neurons (**Figure 3B**).

Of note, the expression of ribosomal protein genes and components of cell growth regulatory pathways, including the mTOR pathway, are dynamically regulated at the mRNA level during the neurodifferentiation process (**Figure 3D-F**). In addition, while many ribosomopathies result in nucleolar stress and p53 activation in other tissues ^12,13^, we did not detect a dramatic increase of p53 levels or induction of a p53-dependent transcriptional response in day 15 mutant organoids relative to the controls (**Figure S5A-C**). This result suggests that the pathology exhibited by *AIRIM* organoids is likely caused by a p53-independent mechanism.

Immunofluorescence staining of control and *AIRIM^V^*^190^*^G^* day 15 organoids confirmed the reduced FABP7/BLBP expression in mutant progenitors (**Figure 4A,B**). Moreover, this analysis revealed that individual rosettes were less well organized in *AIRIM^V^*^190^*^G^* than control organoids. The NE to RG transition is marked by changes in cell morphology^36^. To compare cell shape in control and mutant organoids, we performed sparse GFP labeling, which revealed that NE cells in both control and mutant day 10 organoids appeared wide, columnar, and relatively short (**Figure 4C**). Through days 13 to 15, however, when control progenitors became more elongated, with more constricted apical processes, mutant progenitors remained morphologically similar to those in day 10 organoids (**Figure 4C,D**), consistent with a delay in their RG fate commitment. Together, these scRNA-seq and immunofluorescence data coupled with our proteomic analyses are consistent with a model whereby the *AIRIM^V^*^190^*^G^*mutation causes a temporary delay but not a complete failure in NE to RG differentiation (**Figure 4E**).

**Figure 4.**
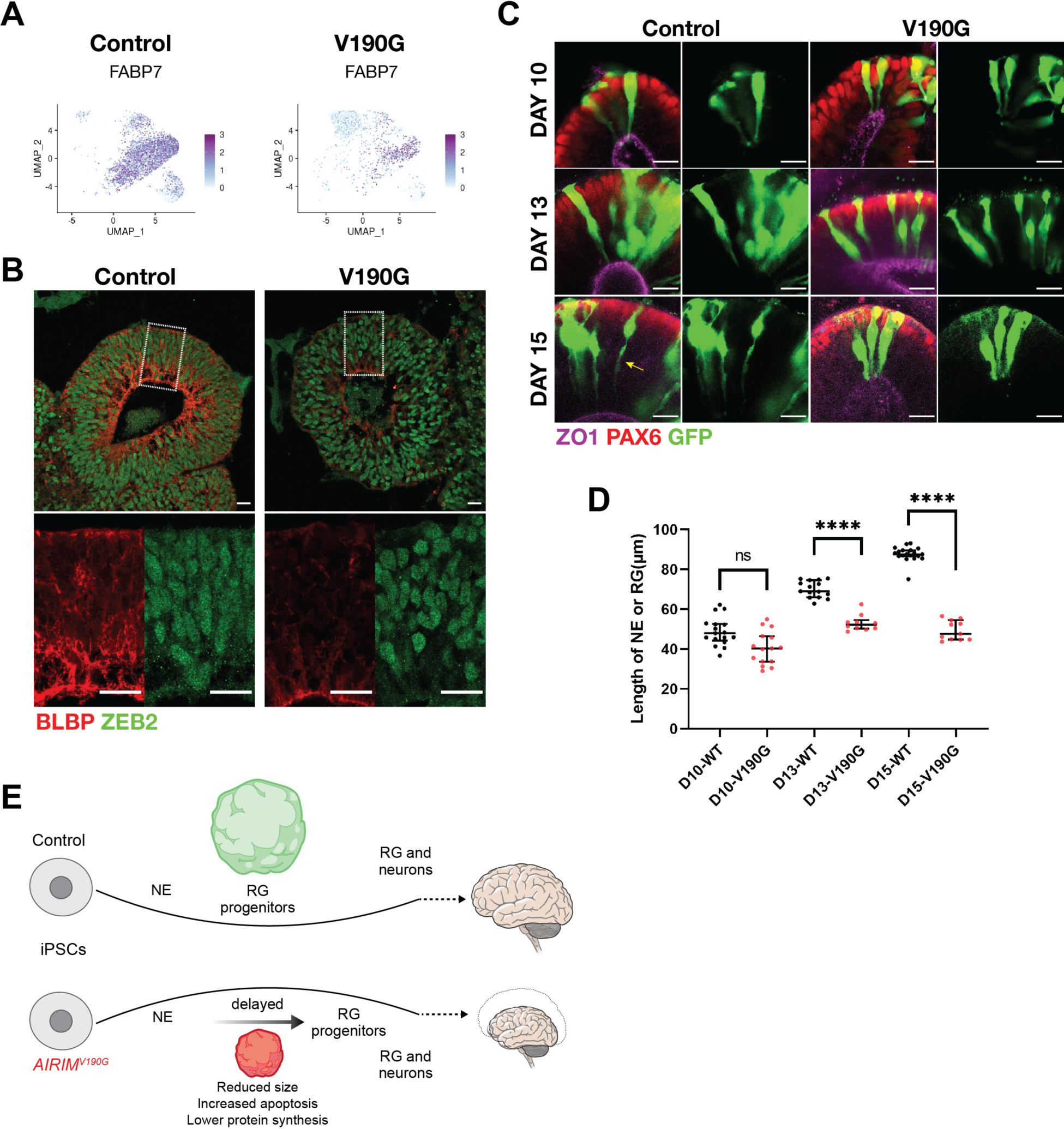
The *AIRIM^V^*^190^*^G^* cerebral organoids exhibit impaired radial glial cell fate specification. (**A**) UMAP showing FABP7 expression in control and AIRIM variant cells. (**B**) Immunofluorescence image of day 15 control and mutant (V190G) organoids of transitioning neuroepithelia marker ZEB2 and committed radial glia marker BLBP (encoded by FABP7). Note that the mutant exhibits less expression of BLBP and less organized nuclei (marked by ZEB2), Scale bar, 50 μm. (**C**) Representative whole mount organoid immunofluorescence images showing the morphology of neural progenitor cells (PAX6+), around apical (ZO1+) lumens, revealed by sparse labeling with GFP in control, and mutant (V190G) organoids. Day 10 cells are columnar and exhibit typical NE morphology. Day 15 control cells show a thinning of apical processes (yellow arrows) while mutant cells still appear columnar. Scale bar, 20 μm. (**D**) Quantification of the length of neural progenitor cells in control and mutant organoids at days 10, 13 and 15, showing significantly reduced length of the progenitor cells in mutant comparing to control organoids. Cells with clear apical and basal labeling were used for quantification. Mann-Whitney U, ∗∗∗∗p < 0.0001, two-tailed, n (day 10 control) = 10 cells, n (day 10 V190G) = 8 cells, n (day 13 control) = 14 cells, n (day 13 V190G) = 16 cells, n (day 15 control) = 8 cells, and n (day 15 V190G) = 8 cells. Error bars are SEM. (**E**) Schematic illustrating observed differences between control and *AIRIM^V^*^190^*^G^* cerebral organoids.

### Decreases in global protein synthesis alters translation of specific transcripts

Translation and ribosome defects first arose on day 10, preceding the observed reduction in cell survival and altered cell fate commitment, as revealed by scRNA-seq (**Figure 2B-E**; **Figure 3**). To further dissect the mechanism underlying the temporal specificity of these phenotypes, we performed ribosome profiling via microfluidic isotachophoresis (Ribo-ITP) ^37^ and assayed ribosome occupancy on mRNAs from single organoids, as well as bulk RNA-seq using day 10 organoids (**Figure 5A; Figure S6A,B; Tables S4, S5**). We observed that a subset of genes important for cell survival, RG commitment, neurogenesis, and early brain development ^38^ exhibited dramatically lower translational efficiency (TE) in *AIRIM^V^*^190^*^G^* organoids relative to the controls, including TPT1, FABP7 and VIM (**Figure 5B-E**). These lower TE genes were also enriched for ribosome components, ribosome biogenesis factors, and other mRNA translation factors. Of note, certain mitochondrial components also exhibited reduced translational efficiency in mutant organoids (**Figure 5C**).

**Figure 5.**
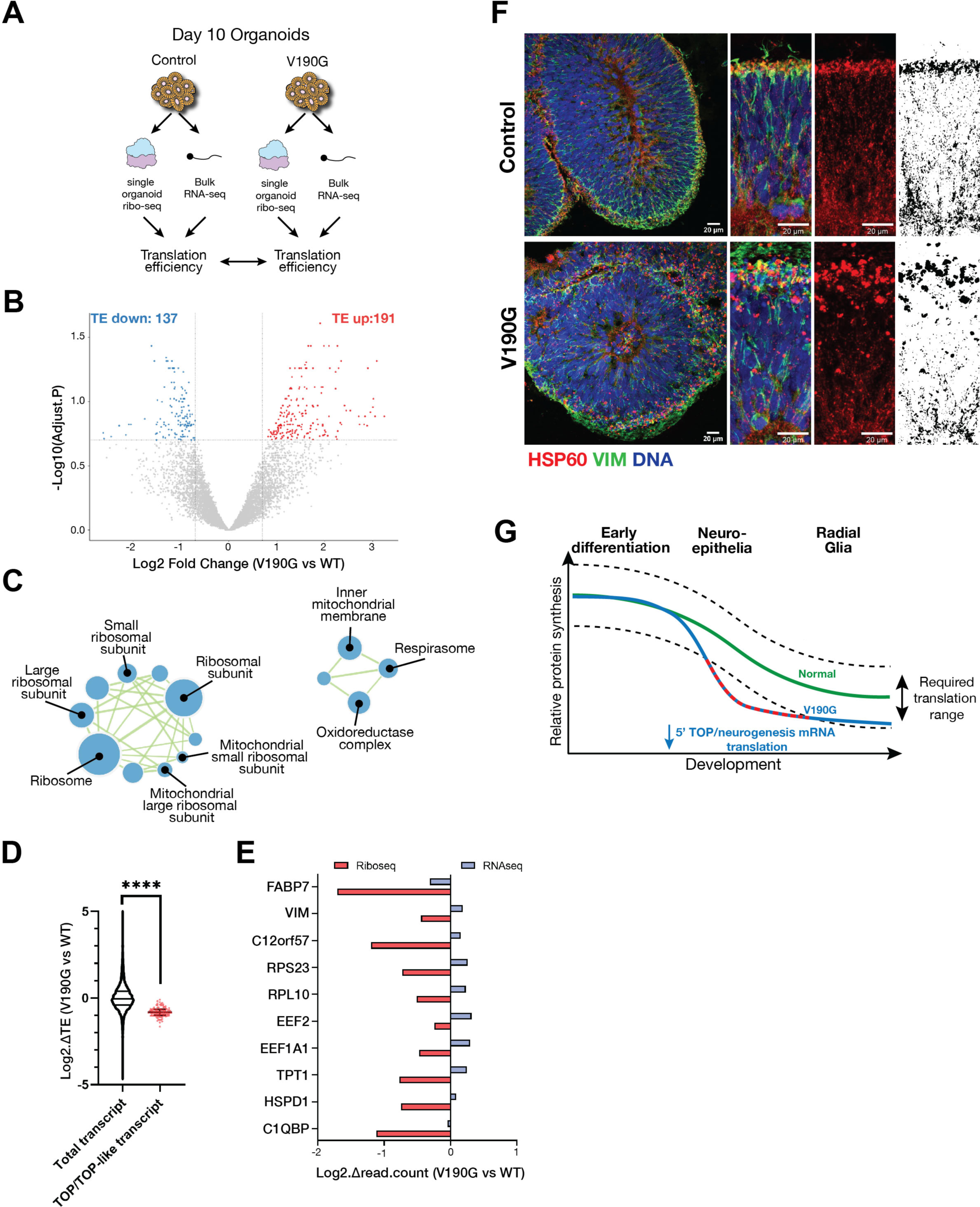
Reduced ribosome availability more profoundly affects a select subset of transcripts in differentiating neuroepithelia. **(A)** Schematic of ribo-seq experiment. (**B**) Volcano plot of change in translational efficiency (TE) of mutant (V190G) relative to control organoids at day 10. red, LFC > 0.7(higher TE in mutant); blue, LFC < -0.7(lower TE in mutant); n = 2 independent batches; FDR < 0.2. (**C**) Gene set enrichment analysis (GO: cellular components) for transcripts exhibit low TE in mutant organoids. (**D**) The TE changes of transcripts whose 5’ UTRs contain 5’ terminal oligopyrimidine (TOP)-like motifs, shown across FDR thresholds for differential translation. ****, p<0.0001, n=12176 total and 105 TOP mRNAs. Unpaired t test with Welch’s correction. (**E**) Relative change in ribo-seq cpm (red) and RNA-seq cpm (blue) of mutant (V190G) day 10 organoids compared to controls for selected high-confidence transcripts. (**F**) Immunofluorescence image of day 15 control and mutant (V190G) organoids of vimentin(green) and mitochondrial matrix HSP60 (red). Note that the mutant exhibits more aggregated mitochondria, Scale bar, 20 μm. (**G**) Model describing how reduced levels of ribosomes in *AIRIM^V^*^190^*^G^* cerebral organoids results in decreased translation of key mRNAs encoding translation machinery components and neurogenesis factors.

Next, we investigated what specific mRNA features contributed to the differential sensitivity to AIRIM complex perturbations and modest reductions in ribosome availability. Previous studies show that the length and structure of 5’UTRs impact mRNA translation^39,40^. Indeed, we found that transcripts with lower TE in mutant organoids showed shorter and less structured 5’UTRs relative to unaffected mRNAs (**Figure S6C,D**). In addition, many of these mRNAs contain 5’ terminal oligopyrimidine (5’ TOP) motifs, which makes them highly sensitive to fluctuations in global translation levels and ribosome availability (**Figure 5D,E**)^40^. TOP/TOP-like elements are found in 83 out of the 137 (61%) transcripts whose translation are disproportionately reduced in mutant organoids. Interestingly, vimentin (VIM), a marker of both RG and the epithelial to mesenchymal transition (EMT), and C12orf57, an important factor for early brain development, contain TOP-like elements within their 5’UTRs ^41–43^. Transcripts of both genes exhibit lower ribosome occupancy in *AIRIM^V^*^190^*^G^* organoids compared to controls (**Figure 5E**). In addition, transcripts encoding the mitochondrial proteins C1QBP and HSPD1 also contain TOP-like elements in their 5’UTR and exhibited reduced translation in mutant organoids (**Figure 5E**). Immunofluorescence analysis of the mitochondrial matrix protein Hsp60 revealed abnormally aggregated mitochondria in day 15 mutant organoids, indicative of impaired mitochondrial function (**Figure 5F**). These data together reveal that a subset of the transcriptome involved in promoting cell survival and neurodevelopment exhibits higher sensitivity to variation in ribosome abundance, thereby conferring vulnerability to mild perturbations in the ribosome biogenesis pathway. Moreover, we speculate that impaired mitochondrial function may contribute to increased cell death in AIRIM organoids.

### Increasing global protein synthesis suppresses neurodevelopmental defects associated with *AIRIM* variants

mTOR signaling controls global protein synthesis levels by phosphorylating key translation factors including eIF4E binding proteins (4E-BPs), resulting in upregulation of cap-dependent translation^44^. mTORC1 is negatively regulated by the TSC1/2 complex ^44^. scRNA-seq revealed expression of mTOR, and other genes that promote global protein synthesis, including Insulin Receptor (INSR) and Lin28, decreased during early neuroepithelial development in human organoids (**Figure 3F**). Concomitantly, expression of TSC1 peaked at day 10 in control organoids. These observations are consistent with OPP labeling and TMT analysis (**Figure 2**). Given that the translation of 5’TOP-element containing mRNAs is highly sensitive to mTOR activity ^40^, we next tested whether increasing mTOR signaling using either genetic or pharmacological approaches could rescue defects associated with the *AIRIM^V^*^190^*^G^* mutation. We hyperactivated mTORC1 by generating heterozygous TSC1 knockout control and mutant iPSCs and used them to generate brain organoids (**Figure 6**). TSC1 haploinsufficiency significantly suppressed the phenotypes associated with *AIRIM^V^*^190^*^G^* mutant organoids, including the organoid growth defects, increased cell death, delayed RG commitment, and aberrant mitochondrial morphology (**Figure 6B-G**).

**Figure 6.**
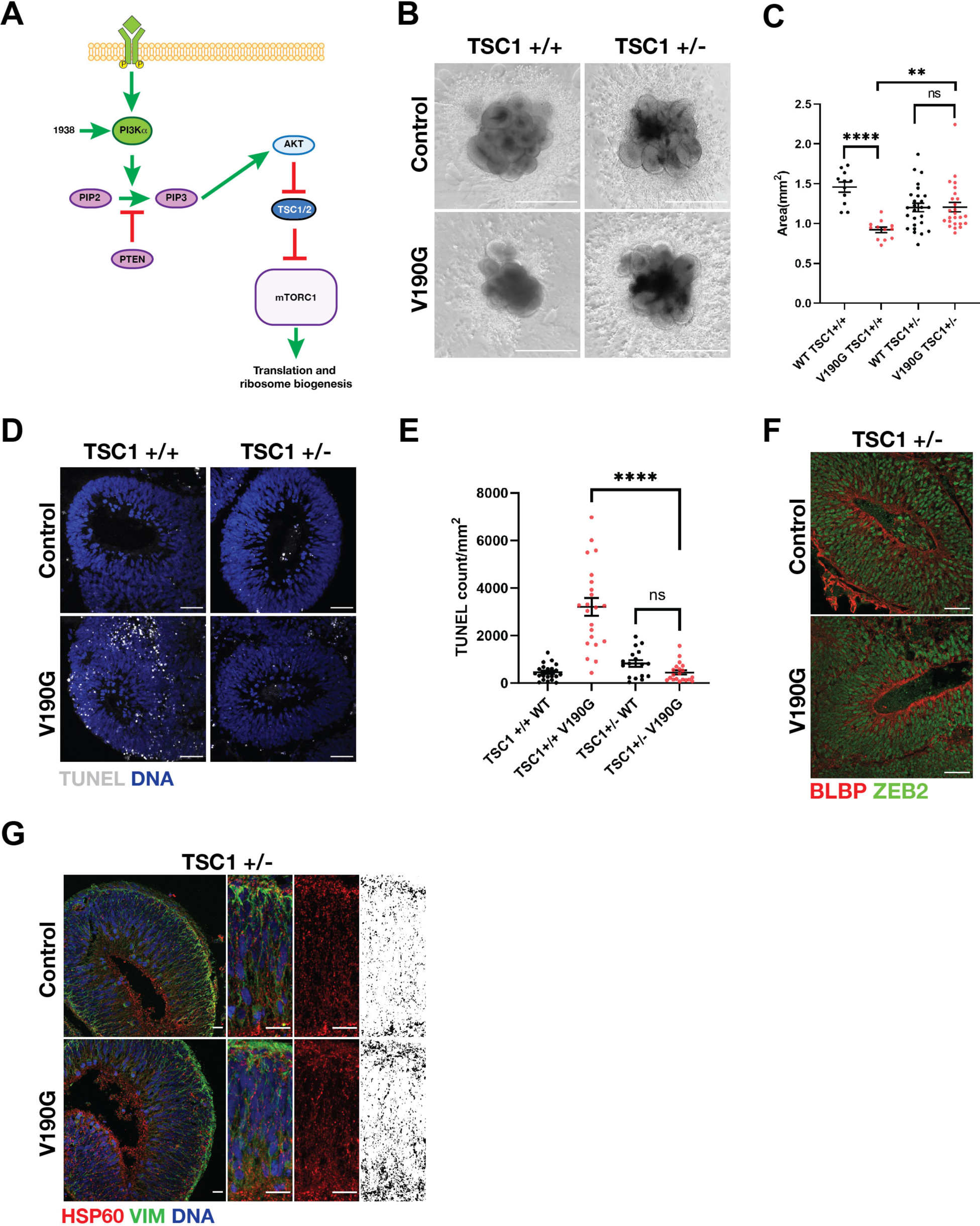
Enhancing global protein synthesis suppresses cell survival and developmental defects within *AIRIM^V^*^190^*^G^* organoids. (**A**) Schematic of mTORC1 mediated regulation of mRNA translation. (**B**) Bright field images of control TSC1+/+, V190G TSC1+/+, control TSC1+/- and V190G TSC1+/- organoids at day 15. Scale bar, 1mm. (**C**) Quantification of bright field images at day 15. V190G TSC1+/- neural tissue is enlarged relative to V190G TSC1+/+; ∗∗p < 0.01; ∗∗∗∗p < 0.0001; Dunettes multiple comparison Unpaired t test with Welch’s correction, n (Day5) = 73 control EBs 62 mutant EBs from 3 independent batches, n (Day10) = 77 control organoids 84 mutant organoids from 4 independent batches, n (Day15) = 46 control organoids 51 mutant organoids from 3 independent batches, error bars are SEM. (**D**) Representative images showing the TUNEL signal in control TSC1+/+, V190G TSC1+/+, control TSC1+/- and V190G TSC1+/- organoids at day 15. Scale bar, 50 μm. (**E**) Quantification of TUNEL signal of individual neuroepithelial bud. TUNEL counts were normalized to the area of the imaged bud. ∗∗∗∗p < 0.0001, Unpaired t test with Welch’s correction. n = 6 control and 6 mutant imaged regions from 2 independent batches. error bars are SEM. (**F**) Immunofluorescence image of day 15 control (WT TSC1+/-) and mutant (V190G TSC1+/-) organoids of transitioning neuroepithelia marker ZEB2 and committed radial glia marker BLBP. Scale bar, 50 μm. (**G**) Immunofluorescence image of day 15 control (WT TSC1+/-) and mutant (V190G TSC1+/-) organoids of vimentin(green) and mitochondrial matrix HSP60(red). Scale bar, 20 μm.

Recent efforts have identified a chemical agonist of PI3Kα, referred to as UCL-TRO-1938, that can be used to increase mTORC1 signaling^45^. We treated *AIRIM^V^*^190^*^G^*organoids with 1 µM of UCL-TRO-1938 starting at day 10 of the differentiation protocol. We assayed control and *AIRIM^V^*^190^*^G^* organoids on day 15 and observed that addition of the compound also alleviated the defects associated with the *AIRIM^V^*^190^*^G^*allelic variant (**Figure 7A-E**). Treatment with UCL-TRO-1938 also rescued the growth and morphological cerebral organoid defects associated with the *AFG2B^I^*^466^*^M/V^*^245^*^E^*variant (**Figure S7**). Together, these results suggest that elevating mTOR signaling counteracts the growth defects and cell fate specification delays associated with AIRIM complex variants in cerebral organoids (**Figure 7E**).

**Figure 7.**
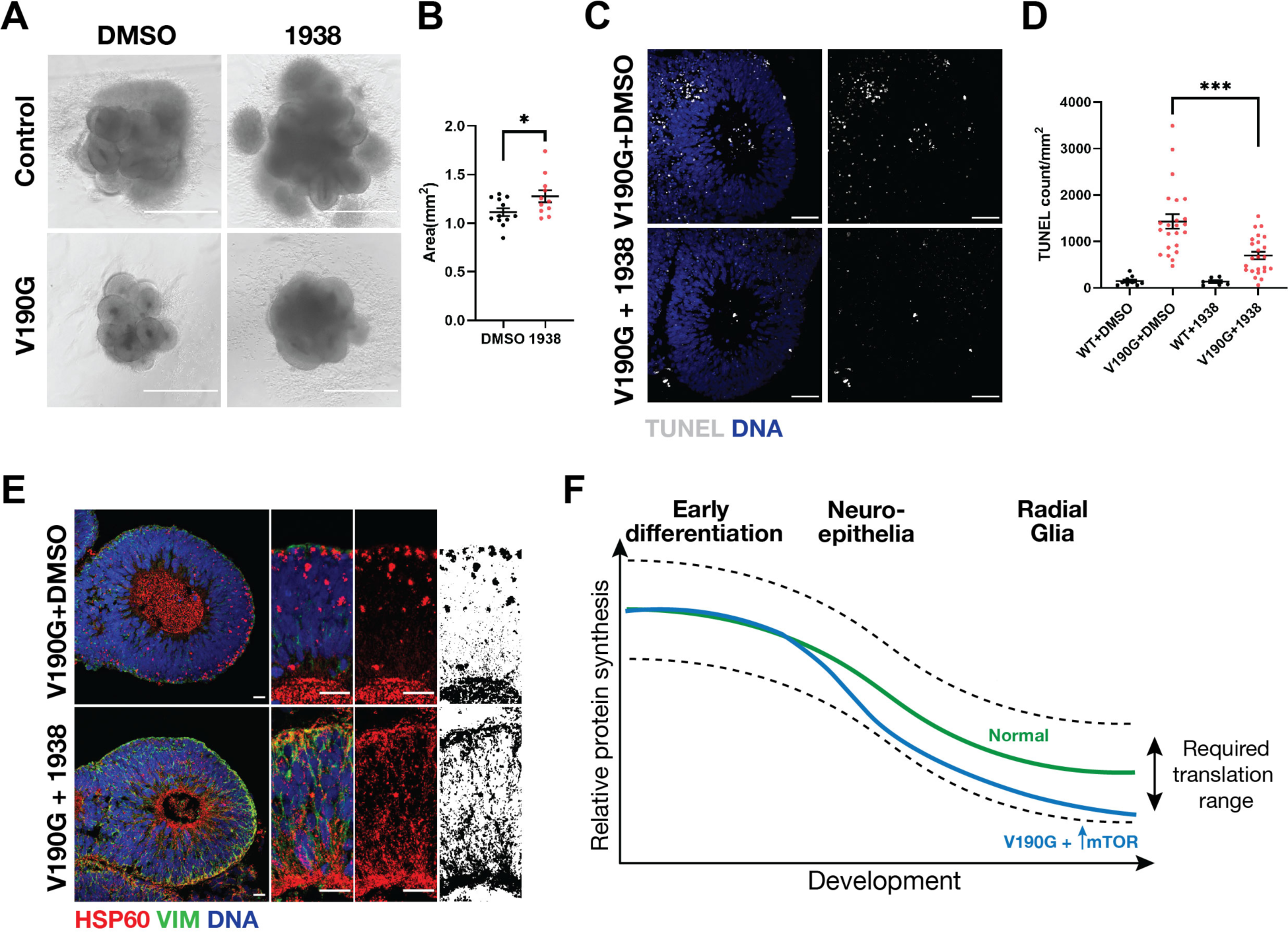
Pharmacological enhancement of mTOR signaling suppresses cell survival and developmental defects within *AIRIM^V^*^190^*^G^* organoids. (**A**) Bright field images of day 15 control (WT) and mutant (V190G) organoids treated with vehicle (DMSO) or 1μM PI3Kα activator UCL-TRO-1938 (1938). Scale bar, 1mm. **(B)** Quantification of size of day 15 mutant(V190G) organoids treated with vehicle (DMSO) or 1μM PI3Kα activator UCL-TRO-1938(1938). ∗p < 0.05. Unpaired t test with Welch’s correction. n=10 organoids from 2 independent batches, error bars are SEM. **(C)** Representative images showing the TUNEL signal in mutant(V190G) organoids treated with vehicle (DMSO) or 1μM PI3Kα activator UCL-TRO-1938(1938) at day 15. Scale bar, 50 μm. **(D)** Quantification of TUNEL signal of individual neuroepithelial bud. TUNEL counts were normalized to the area of the imaged bud. ∗∗p < 0.01, Dunnett’s T3 multiple comparisons test. n (WT+DMSO) = 9, n(V190G+DMSO)=11, n(WT+1039)=7, n(V190G+1938)=13 imaged regions from 2 independent batches. error bars are SEM. (**E**) Immunofluorescence images of day 15 control (V190G, DMSO 0.1%) and treated (V190G, UCL-TRO-1938 1μM) organoids stained for vimentin (green) and mitochondrial matrix HSP60 (red). Scale bar, 20 μm. (**F**) Model describing a potential explanation for the observed suppression of AIRIMV190G phenotypes by UCL-TRO-1938.

## Discussion

Collectively, our data support a model whereby the dynamic regulation of global mRNA translation plays a significant role in early brain development. Leveraging iPSC-derived cerebral organoids, our study identifies a critical stage of early human brain development during which ribosome numbers decline but global protein synthesis must be maintained above a minimum threshold to support cell survival and cell fate specification. Previous studies have shown the importance of the neuroepithelial to radial glial cell transition in regulating the brain size across different primate species^36^. Stage specific variation in ribosome availability may render differentiating cells sensitive to minor perturbations in ribosome biogenesis.

Here, we identify 5 *AIRIM* allelic variants across seven different families that are tightly associated with a range of neurodevelopmental disorders. These phenotypes are strikingly similar those associated with alleles in other components of the AIRIM complex, including AFG2A, AFG2B, and CINP^19–22,24,25,31^ and are predominantly associated with global developmental delay, intellectual disability, muscular hypotonia, limb spasticity, dystonia, microcephaly, infantile seizures, as well as hearing and vision impairment (see **Figure S1E** for comparative summary). Importantly, patients that carry *AFG2A* and *AFG2B* allelic variants commonly exhibit similar neuroimaging features ^21–24^. It is worth noting that the severity of the imaging findings, particularly the extensive atrophy, appears slightly more pronounced in *AIRIM* patients compared to those described for *AFG2A*. Certain pathogenic variants, such as those in *UBTF*, can result in abnormal ribosomal RNA (rRNA) expression, leading to severe neuroregression and a similar neuroimaging phenotype characterized by prominent supratentorial brain atrophy with thinning corpus callosum, abnormal myelination, with relative sparing of the cerebellum and brainstem ^46^. Mutations in tRNA production machinery have also been associated with NDDs ^47^. Perhaps these defects converge on a common mechanism, such as inappropriate declines in mRNA translation during specific stages of brain development when ribosome availability or activity may be limited.

We find allelic variants in both *AIRIM* and *AFG2B* result in widespread cell death and delays in RG specification within the context of cerebral organoids. We speculate that cells undergoing the NE to RG transition need to maintain a minimum threshold of protein synthesis capacity to ensure proper differentiation in a timely manner. Unexpectedly, *AIRIM* and *AFG2B* variants did not cause obvious widespread activation of a p53-dependent stress response. As cells from patients carrying variants in other components of the AIRIM complex exhibit mitochondria defects^22^, we speculate that the increased cell death observed in mutant organoids is most likely caused by mitochondrial malfunction and clumping. Thus, our results provide a unified molecular framework that potentially links defects in cytoplasmic ribosomes with mitochondrial function.

Our study further reveals that reductions in protein synthesis capacity affect the translation of specific mRNAs in developing brain organoids. Most of these mRNAs contain 5’ TOP elements. Recent work has found that transcriptional start sites in individual genes can vary across tissues, resulting in the cell-type specific inclusion of 5’ TOP and TOP-like elements with the 5’UTRs of mRNAs^48^. Our results indicate that minor perturbations in 60S maturation become exacerbated during neurodifferentiation through reductions in the translation of mRNAs involved in protein synthesis. Increasing mTOR activity partially suppresses the *AIRIM* variant phenotypes, further linking the abnormalities observed in patients with defects in the regulation of protein synthesis. However, ribosome and global protein synthesis levels must be finely tuned during different developmental events. Inappropriate increases in global protein production can also have deleterious effects on brain development. For example, mutations that disrupt the TSC1/2 complex result in non-cancer tumor growth in human brain^44^. Therefore, ribosome availability and protein synthesis will need to be adjusted in space and time to successfully treat abnormalities caused by loss of function mutations in ribosome assembly machinery. These results encourage future investigation into therapeutic approaches aimed at modulating protein synthesis in a cell-type and temporally specific manner.

## Materials and Methods

### Patient recruitment and sequencing

Patients were recruited as part of large-scale sequencing screens as previously described ^24^.Exome sequencing was performed in a research setting or in accredited molecular diagnostic laboratories. Sanger sequencing was used for variant validation and segregation analysis. Variants were mapped to the predicted structure of AIRIM_49,50._

### Stem cell culture and proliferation assay

Five human iPSC lines (SCVI274, F856/parent#1, F856/parent#2, F740/proband#1, F740/proband#2) were used in this study. SCVI274 were obtained from Stanford SCVI BioBank. F856#1, 2 and F740#1, 2 were reprogramed in this study. All iPSC lines were cultured in mTESR-plus medium(Stem Cell Technologies, 100-0276) on matrigel-coated plates(Corning, 354277). Cells were passaged every 3-5 days after dissociation with ReLeSR(Stem Cell Technologies, 100-0483). The medium was supplemented with Rho-associated protein kinase (ROCK) inhibitor Y-27632(Tocris, 1254) at a final concentration of 10 µM for the first 24h after passaging. Routine mycoplasma tests were performed with Universal Mycoplasma Detection Kit(ATCC, 30-1012K). For proliferation assay, cells were dissociated with ACCUTASE(Stem Cell Technologies, 07920) and were passaged onto Matrigel-coated 12-well plates at 1.0×105 cells/well in mTESR-plus medium with 10 µM ROCK inhibitor. 24h after passage, medium was changed with mTESR-plus without ROCK inhibitor. Medium was then changed every 24h. Cells were dissociated with TrypLE Express Enzyme(ThermoFisher, 12605010) and were counted by hemocytometer.

### Reprogramming of fibroblasts

The fibroblasts were reprogrammed using episomal vectors as previously described^51^. Briefly, 1×106 fibroblasts were electroporated with 2.5 µg of each of the following vectors: pCXLE-EGFP, pCXLE-hOCT3/4-shp53-F, pCXLE-hSK, and pCXLE-hUL (Addgene #27082, #27077, #27078, #27080, respectively) using a NEPA21 Type II Super Electroporator (Bulldog-Bio) using the manufacturer’s recommended parameters. After electroporation, fibroblasts were collected and transferred equally to a 6-well plate coated with Matrigel in DMEM (Sigma) supplemented with 10% fetal bovine serum (FBS). After 48 hours, medium was replaced with mTESR-plus medium. Medium was refreshed every 24 hours from then on. After 3-5 weeks, tentative human iPSC colonies were manually isolated and expanded for analysis.

### CRISPR-mediated gene editing and knock out

For the SCVI274 AIRIM mutant line, 2×106 parental cells were resuspended in 100uL OPTI-MEM with 5µg of px458-sgRNA plasmid and 5µg of ssDNA donor. The cells were then electroporated at poring pulse 150V/length 2.5ms/interval 50ms, transfer pulse 20V/length 50ms/interval 50ms. 48h after electroporation, cells were sorted for GFP positive on a FACS-Aria sorter. The sorted cells were plated onto Matrigel coated 6-well plates at 2,000 cells per well in mTESR-Plus with 10 µM ROCK inhibitor. 7-14 days after plating, colonies were picked into 24-well plate. Colonies were screened by PCR and Sanger sequencing. Homozygous mutant clones were expanded, and genotypes were reconfirmed after expansion. For the TSC1 knock out, 2×106 parental cells were resuspended in 100uL OPTI-MEM with 5µg of px458-sgRNA plasmid. The cells were electroporated, sorted and screened as described above. Sequences of sgRNA, donor and PCR primers are available in supplementary table S5.

### Teratoma assay

hiPSCs were dissociated into a single-cell suspension, then resuspended at a concentration of 1×107 cells/mL in a mixture of 50% Matrigel and 50% culture medium containing ROCK inhibitor. Cells were injected subcutaneously into the flanks of female NOD/SCID immunodeficient mice with a total of 1×106 cells per injection site. Teratomas were isolated after 8 weeks of growth once visible tumors had formed, and the tissue was fixed in 4% PFA for 48 hours. Tumor samples were submitted to the UT Southwestern Histopathology Core facility for paraffin embedding, sectioning, and hematoxylin/eosin (H&E) staining. Germ layer contribution was determined through imaging and histologic examination of H&E-stained tumor sections.

### Generating cerebral organoids

Cerebral organoids were generated using Stemdiff Cerebral Organoid Kit (Stem Cell Technologies, 08570). The timing of the kit protocol (StemCell Technologies Document #DX21849) was followed. The EB formation step was modified in order to generate organoids with enhanced telencephalic identity ^36^. In Aggrewell 800 plates (Stem Cell Technologies, 34815), 6×105 cells in 2mL EB formation medium were added per well to achieve 2000 cells per EB. During the EB formation period, the medium was changed 50% every day with an electronic auto dispenser at slowest pipetting speed. At neural induction, Aggrewell were changed with induction medium for 3 times in order to achieve <0.5% remaining EB formation medium. Matrigel embedding for individual organoids was described in (Lancaster Nature protocol). Matrigel embedding for large-scale assay (>100 organoids/batch) was performed by following the previously described protocol ^36^. All comparisons between samples and treatments were performed on organoids generated using identical protocols. For the sparse labeling of neural progenitor cells, 5 mL CytoTune emGFP Sendai fluorescence reporter (ThermoFisher, A16519) was added to Aggrewells when the organoids were switched to neural induction medium.

### Organoid size analysis

Images of organoids in culture were taken with an inverted microscope(ECHO R4) at x4 magnification objective. Area values were obtained by quantifying individual organoids on ImageJ, which measured area in mm^2^.

### Immunofluorescence

For cryosectioning, organoids were washed in PBS before fixing in 4% PFA at 4 °C overnight. After fixation, organoids were washed three times with PBS and were then incubated in PBS/15% sucrose for 6h at 4 °C, and then PBS/30% sucrose for 16-24h at 4 °C. Organoids were transferred to plastic cryomolds and were embedded in OCT compound for snap-freezing on dry-ice and were cryo-sectioned at a thickness is 20µm. Sections were warmed at RT for 10min and rinsed with TBS for three times with 10min each before incubating with blocking solution (0.3% Triton X-100, 5% donkey serum in TBS). Sections were then incubated with primary antibodies in blocking solution for 16-24h at 4 °C. After primary antibody, sections were washed with TBS for three times with 15min each before incubating with secondary antibodies in blocking solution for 2h at RT. Sections were then washed three times with TBS with 15min each. Sections were then incubated with Hoechst 33342 at 2µg/mL for a 15min incubation at RT. Slides were mounted with ProLong Gold Antifade Mountant (Thermofisher, P36930). Stained organoid cryosections were imaged using a ZEISS LSM 800 confocal microscope, at 0.7-0.8µm intervals using a x40 magnification objective. The images were further processed using Fiji. For whole mount imaging, organoids were washed and fixed the same as cryosection. Organoids were then permeabilized in blocking solution for 16-24h at 4 °C. After permeabilization, organoids were incubated with primary antibody in blocking solution for 16-24h at 4 °C. Organoids were then washed in PBS/0.04% BSA for three times with 15min each. Organoids were then incubated with secondary antibodies in blocking solution for 2h at RT. Optionally, Hoechst 33342 was added into secondary antibody incubation at 2µg/mL. Organoids were then washed three times with PBS/0.04% BSA with 15min each and were then stored in PBS at 4 °C before imaging. Stained organoid cryosections were imaged using a ZEISS LSM 780 confocal microscope, at 8-12µm intervals at x20 magnification objective. The images were further processed using Fiji.

### TUNEL assay

Cryosectioning for TUNEL assay is the same as previously described in immunofluorescence. Sections were warmed up, washed with PBS two times in for 5 min each and were then permeabilized in PBS/0.2% Triton/0.5%BSA for 30min. After permeabilization, sections were washed with PBS two times in for 5 min each. TUNEL assay reaction was carried out with TUNEL Assay Kit Fluorescence, 594 nm (CellSignaling, 48513) according to the manufacturer’s directions.

### OP-Puro-click

One day before labeling, medium of iPSCs, EBs and organoids was changed to ensure nutrient level. On the day of labeling, the culture was changed with medium containing 20µM O-propargyl-puromycin (OPP, Clickchemistrytools, 1407). Samples were incubated at 37 °C for 30min and were then processed according to the downstream assay. For whole mount sample, EBs were fixed as described in immunofluorescence. After fixation, EBs were permeabilized in DPBS/0.25% Triton X-100 for 20min and were washed once with DPBS/3% BSA before click reaction. For cryosectioned samples, sections were warmed up, washed with DPBS for 3 times 5min each, permeabilized in DPBS/0.25% Triton X-100 for 20min and were washed once with DPBS/3% BSA before click reaction. Click reaction was carried out with Click-&-Go Cell Reaction Buffer Kit (Clickchemistrytools, 1263) with AZDye 568 Azide (Clickchemistrytools, 1291) according to the manufacturer’s directions.

### Tandem mass tag (TMT) mass spectrometry

For protein lysate, iPSCs, EBs and organoids were scraped or collected directly by pelleting at 500g x5min. Samples were washed once with DPBS, and the pellet were snap-frozen at -80 °C until lysis. Samples were lysed in RIPA buffer (ThermoFisher, 89900) with 0.5µL/mL Benzonase (Sigma, E1014) for 15min on ice. Lysate was centrifuged at 15000xg for 15min at 4 °C. The supernatant was collected and submitted to UTSW Proteomics Core for TMT-18plex. Data was analyzed using Proteome Discoverer 3.0 and was searched using the human protein database from UniProt.

### Dissociation of brain organoids and scRNA-seq

Organoids were dissociated as previously described^52^. In brief, iPSCs and EBs were dissociated in TryplE (ThermoFisher, 12605010). Organoids on day10, 15 and 30 were dissociated in neural dissociation kit-P (miltenyibiotec, 130-092-628). scRNA-seq libraries were generated using the Chromium Single Cell 3′ Library & Gel Bead Kit v3 or v3.1 (10x Genomics, PN-1000075, PN-1000121). For each sample, an estimated 10,000 cells were loaded. Libraries were sequenced on a N2K2.

### scRNA-seq data processing

Reads from scRNA-seq were aligned to the GRCh38 human reference genome and the cell-by-gene count matrices were produced using the Cell Ranger pipeline (10x Genomics). Data were analyzed using the Seurat R package v.3.1.5 using R v.4.2. Data from 10 individual runs were merged. Cells were filtered on the basis of unique molecular identifier (UMI) counts(>500), the number of detected genes(>500), the log10genes per UMI(>0.8), and the fraction of mitochondrial genes(<0.2). UMI counts were normalized for each cell by NormalizeData() with a scale factor of 10,000. We selected the union of the 100 most variable genes for each time point separately (local) as well as across the full dataset (global). The local and global sets were then combined with a previously reported organizer set^35^. Cell-cycle related genes were excluded from the set based on AnnotationHub. The data were then scaled and the cell cycle scores and mitochondrial scores were regressed out by ScaleData(). Principal Component Analysis (PCA) was performed using the Seurat function RunPCA(). The first 10 PCs were used to integrate the different time points and genotypes in the dataset using Harmony with max iteration as 50. We performed UMAP using RunUMAP() with reduction method as “harmony” and otherwise default settings. Cells were clustered in PCA space using FindNeighbors(), followed by FindClusters with resolution = 0.8. Differentially regulated genes in each cluster were identified using FindAllMarkers() with logFC thereshold at 0.25. The top significant differentially regulated genes were selected for heatmap visualization. Features used for integration can be found in supplementary table S6. Ribosome biogenesis gene set used in analysis are listed in supplementary table S7.

### Ribo-seq sample preparation and data analysis

Single wild type and *AIRIM^V^*^190^*^G^*brain organoids were collected in a 0.5ml tube and subjected to cell lysis by freeze and thaw in the presence of Ribo-ITP lysis buffer. The resulting homogenized cells were then treated with RNase I at 37°C for 15 minutes, and the reaction was stopped by adding 2% SDS. The remaining steps of Ribo-ITP experiments were performed as previously described ^37^. Ribosome profiling sequencing libraries were prepared using the D-Plex Small RNA-seq kit (C05030001, Diagenode) with slight modifications. The dephosphorylation reaction was supplemented with 0.5 μl T4 PNK (NEB), and the reaction was incubated for 25 minutes. Subsequently, the complementary DNA (cDNA) was amplified for 12 PCR cycles. We used AMPure XP bead cleanup (1.8X), followed by size selection using 3% agarose, dye-free gel cassettes with internal standards (BDQ3010, Sage Science) on the BluePippin platform. Sequencing was performed using Illumina Novaseq 6000.

Ribosome profiling data were processed using RiboFlow with the following modifications ^53^. Unique molecular indices (UMIs) corresponding to the first 12 nucleotides from the 5’ end of each read was extracted using UMI-tools ^54^. The four nucleotides following the UMI were discarded as they are incorporated during the template-switching reverse transcription step. Transcriptome-mapping reads with mapping quality greater than two were retained using the default settings in RiboFlow. PCR duplicates were removed using UMIs. Finally, ribo files were generated using Riboflow, and further analyses, including metagene plots and quality control were generated using RiboR ^53^.

### Cytoplasmic RNA-seq sample preparation and data analysis

Organoids were produced with bulk-embedding as described above (Generating cerebral organoids). On day10, 300x organoids were collected and were dissociated from matrigel by centrifuging at 500xg for 5min. To extract cytoplasmic RNA, organoid pellet was lysed in lysis buffer (20mM Tris-Cl pH7.5, 5mM MgCl_2_, 150mM NaCl, 1mM DTT, 0.1% NP-40). Lysate was collected into 1.5mL tube and incubated on ice for 15min. Lysate was then centrifuged at 21130g at 4°C for 10min to remove nuclei, mitochondria and debris. The supernatant was collected and the cytoplasmic RNA was extracted by Trizol-LS (ThermoFisher, 10296028) following the manufacturer’s protocol. The strand specific whole transcriptome (ribo-minus) sequencing was performed at The McDermott Center Next Generation Sequencing (NGS) Core at UT Southwestern using TruSeq Stranded mRNA Sample Preparation Kit (Illumina). The reads were then used as reference of single organoid ribo-seq under same condition. The 5’ adapter sequence ‘AGATCGGAAGAGCACACGTCTGAACTCCAGTCA’ was clipped from the first read, and processed using RiboFlow to quantify read counts reads per transcript.

### Differential translation efficiency analysis

Using RiboR, we extracted read counts corresponding to coding regions from the ‘ribo’ files across all experiments. All analyses utilized ribosome footprints ranging between 26 and 33 nucleotides in length. We performed a joint analysis and normalization of ribosome occupancy and RNA-seq data employing the TMM normalization method ^55^. We then calculated transcript-specific dispersion estimates and assessed differential translation efficiency using the edgeR tool ^56^. Specifically, to identify genes with differential translation efficiency, we evaluated changes in ribosome profiling while adjusting for differences in RNA expression using a generalized linear model, in which we considered RNA expression and ribosome occupancy as two experimental manipulations of the cell’s RNA pool ^57^. We employed the Benjamini–Hochberg procedure to compute adjusted p-values.

## Statistics and reproducibility

### Organoid size analysis

For AIRIM across time points. For Day5: *n* = 73 for total control EBs, *n* = 62 for total mutant EBs, from 3 experimental batches. For Day10: *n* = 77 for total control organoids, *n* = 84 for total mutant organoids, from 4 experimental batches. For Day15: *n* = 46 for total control organoids, *n* = 51 for total mutant organoids, from 4 experimental batches. For Day30: *n* = 12 for total control organoids, *n* = 12 for total mutant organoids, from 2 experimental batches. *P* values were calculated using unpaired t test with Welch’s correction. For TSC1: n=11 for total control TSC1+/+ organoids, n=12 for total V190G TSC1+/+ organoids, n=37 for total control TSC1 +/- organoids, n=35 for total V190G TSC1+/- organoids, from 2 experimental batches. *P* values were calculated using Dunnett’s T3 multiple comparisons test.

### Proteomic analysis

For each batch, 300 EBs, 300 day10 organoids and 100 day15 organoids were used for control and V190G mutant. Samples from three individual batches were analyzed. To detect the abundance of ribosomal proteins across conditions, RM one-way ANOVA and Tukey’s multiple comparison were performed.

### scRNA-seq analysis

At least 50 day 5 embryoid bodies, 3-6 day 10 neuroepithelia, 3-6 day 15 organoids and 3 day 30 organoids of each genotype were pooled for each dissociation. For each condition, approximately 10,000 cells were sequenced.

### Single organoid ribo-seq analysis

2 control and 3 mutant organoids on day 10 were sequenced.

### Bulk RNA-seq analysis

300 organoids were pooled and sequenced per genotype, per replicate, with a total of 2 biological replicates.

### Immunofluorescence/TUNEL/OP-Puro imaging

At least 12 organoids of each condition were used for the assays. P values were calculated from Welch’s t test(two conditions) or Kruskai-Wallis test(multiple comparison).

## Acknowledgments

We acknowledge the UTSW Flow Cytometry Facility and Moody Foundation Flow Cytometry Facility at UT Southwestern for performing cell sorting/flow cytometry analysis for this project. We acknowledge the Next Generation Sequencing Core of McDermott Center at UT Southwestern for performing next generation sequencing for this study. We thank the UTSW Medicinal Chemistry Core and Monika Antczak at UT Southwestern for performing synthesis of small molecules for this study. We acknowledge UTSW Proteomics Core at UT Southwestern for proteomic analysis for this study. We thank J. Mendell, K. Orth, M. Sieber, and B. Ohlstein for discussion, suggestions and critically reviewing the manuscript. The authors would like to thank the families for participating in this study.

## Author contributions

Conceptualization: CN, LY, JW, MB

Methodology: CN, LY, BV, DAS, EB, YLW, YD, YL, AK, CX, CC, RM

Investigation: CN, LY, BV, DP, H-GK, CE, EGK, SI, FE, RSB, EMCS, MD, MK, MYVM, ZM, HE, EK, SMNA, AC, CAPFA, HH, FSA, RM

Visualization: CN, YLW, YW, MS, YD

Funding acquisition: BV, CC, DAS, JW, MB

Project administration: RM, JW, MB Supervision: RM, CC, JW, MB

Writing – original draft: CN, MB, BV, RM

Writing – review & editing: CN, MB, JW, LY, BV, RM

## Data and materials availability

sc-RNA-seq and ribo-seq data can be found GEO####. All other data are available in the main text or the supplementary materials.

## Supplementary Materials

Figures S1-S7

## Other Supplementary Materials

Tables S1 to S8

