## Supplemental Figures S1-S7 for "An inappropriate decline in ribosome levels drives a diverse set of neurodevelopmental disorders"

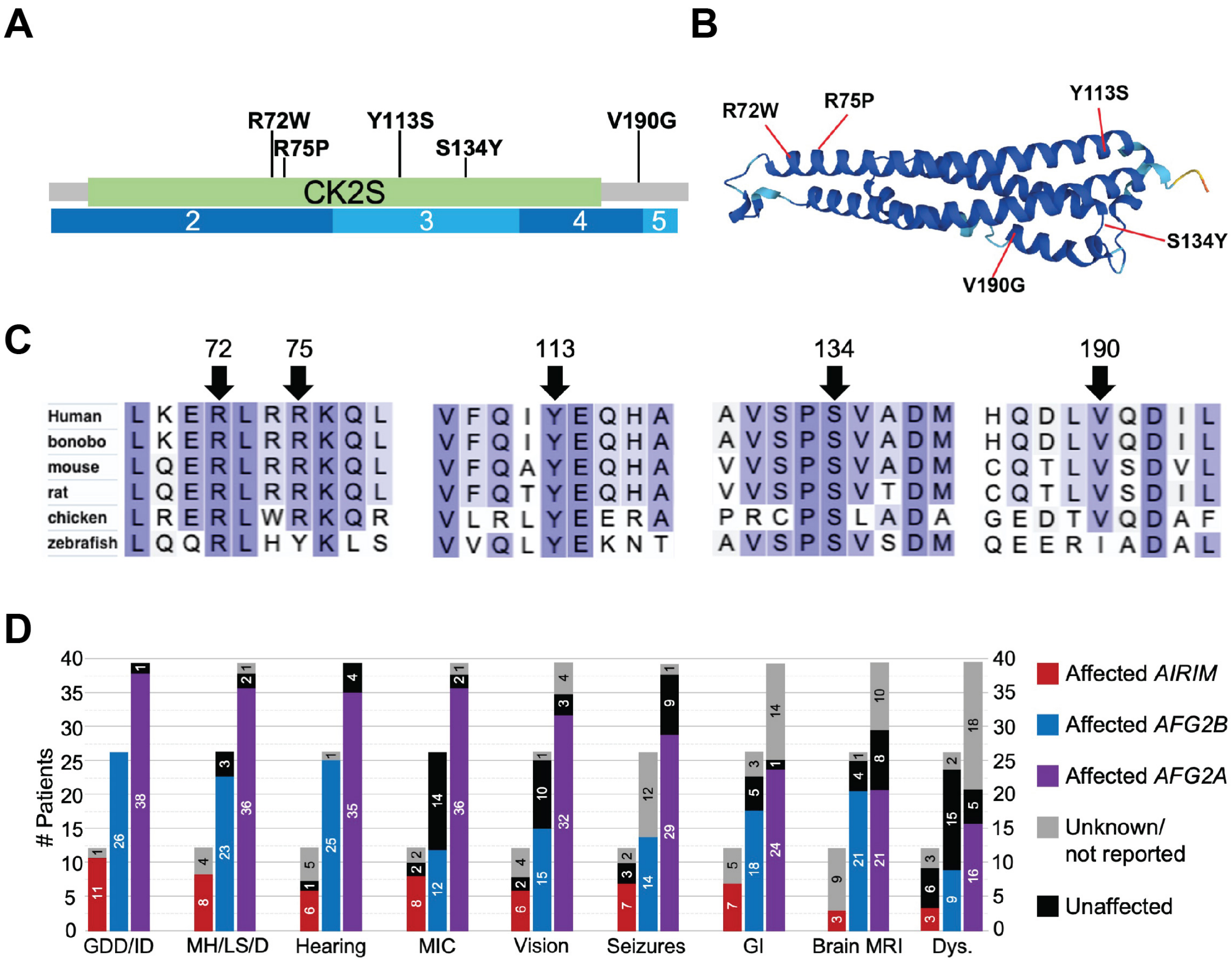


**Figure S1. Association of *AIRIM* variants with neurodevelopmental disorders.** (**A**) Schematic of the protein with variants marked. The casein kinase II substrate (CK2S) shown in green with the corresponding AIRIM exons in blue below. (**B**) Mapping of variants onto the predicted structure of AIRIM. (**C**) Alignment of multiple AIRIM orthologues with amino acid substitutions marked with arrows. (**D**) Bar graphs summarizing the comparative proportions of various clinical findings in individuals with *AIRIM*, *AFG2B* and *AFG2A* variants.

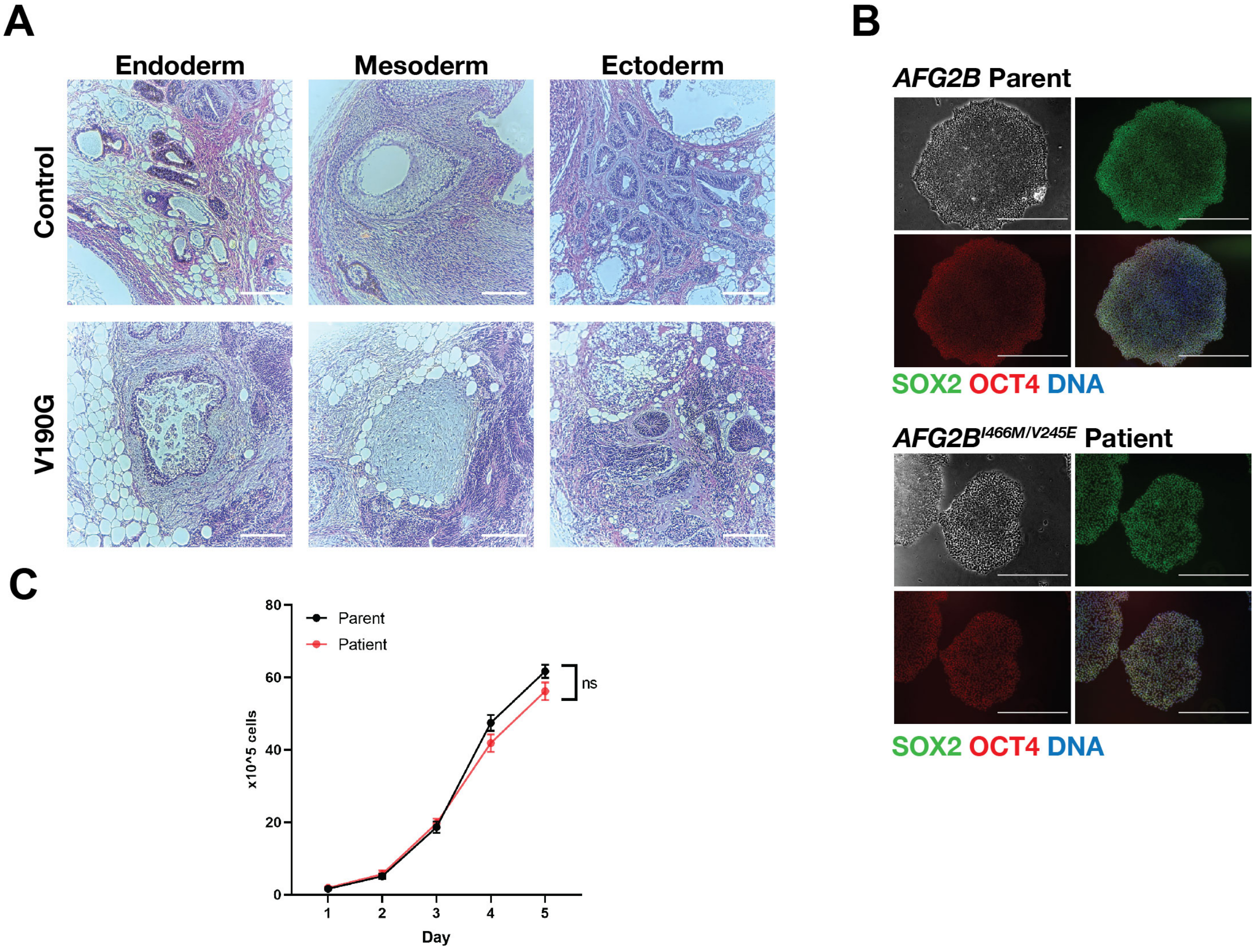


**Fig. S2. Proliferation and pluripotency assays of iPSC lines. (A)** Teratoma assay of SCVI274(control) and the derived *AIRIM*$^{V190G}$ mutant(V190G) lines. Scale bar, 200 µm. **(B)** Immunofluorescence image of pluripotency markers SOX2(green) and OCT4(red) of iPSC clones derived from the AFG2B patient and parental control fibroblasts. Scale bar, 1mm. (**C**) Growth of parental and *AFG2B* patient lines. Two-way RM ANOVA. n=4 individual wells quantified per genotype, per time point.

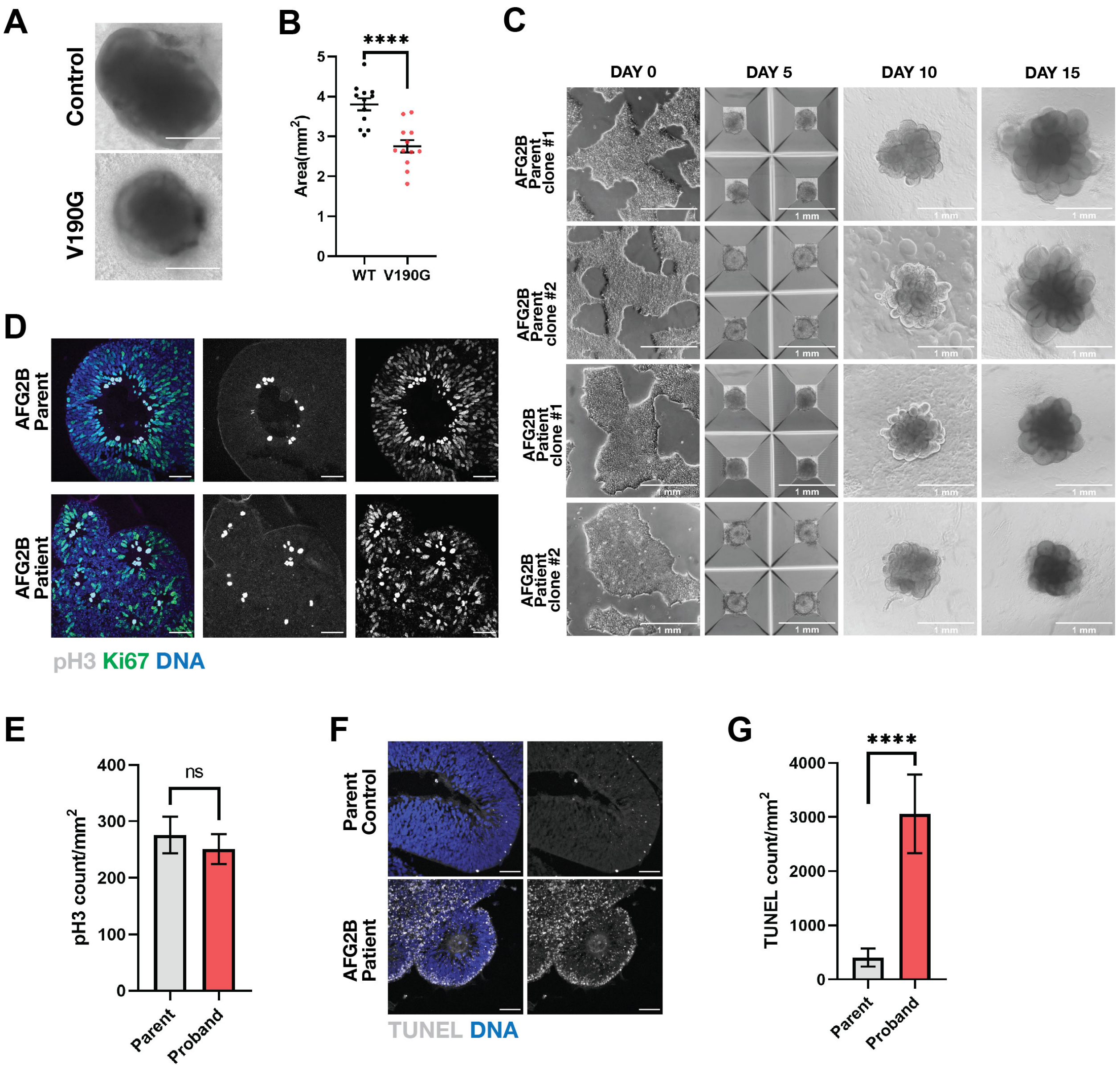


**Fig. S3. A disease associated AFG2B variant induces stage-specific growth defects of cerebral organoids. (A)** Bright field images of SCVI274(control) and the *AIRIM*$^{V190G}$ mutant(V190G) organoids at day 30. Note both control and mutant organoids show cortical plate structure. Scale bar, 1mm. (**B**) Quantification of bright field images at day 30. show Control neural tissue is enlarged relative to mutant(V190G); ****p < 0.0001, Unpaired t test with Welch's correction, n= 12 control organoids and 12 mutant organoids from 2 independent batches error bars are SEM. **(C)** Parental control and AFG2B patient derived EBs and organoids at days 5, 10, and 15. Scale bar, 1mm. Note organoids from patient derived iPSCs exhibit reduced growth and less elongated neuroepithelial buds on day 15. (**D**) Representative images showing the proliferation marker phospho-Histone 3(pH3, grey) and Ki67(green) signal in control (parent) and mutant (AFG2B patient) day 15 organoids. Scale bar, 50 µm. **(E)** Quantification of pH3 signal of individual neuroepithelial bud at day 15 control(parent) and mutant (AFG2B patient) organoids. Unpaired t test with Welch's correction. n = 10 control and 14 mutant imaged regions from 2 independent batches. error bars are SEM. **(F)** Representative images showing the TUNEL signal in control(parent) and mutant (AFG2B patient) day 15 organoids. Note mutant organoids exhibit significantly more TUNEL signal. Scale bar, 50 µm. **(G)** Quantification of TUNEL signal of individual neuroepithelial bud showing mutant organoids exhibit more TUNEL signal. TUNEL counts were normalized to the area of the imaged bud. ****$p < 0.0001$, Unpaired t test with Welch's correction. n = 10 control and 11 mutant imaged regions from 2 independent batches. error bars are SEM.

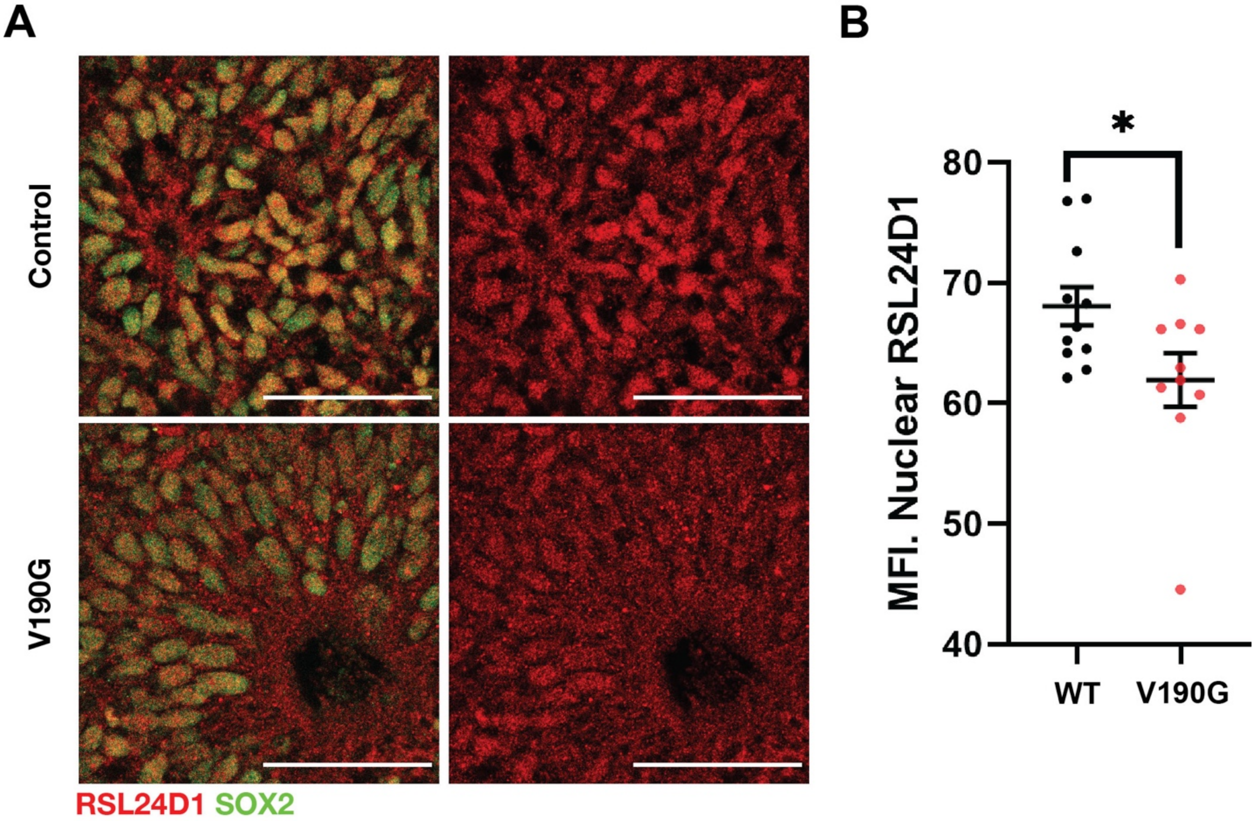


**Figure S4 *AIRIM*$^{V190G}$ variant organoids exhibit mis-localization of RSL24D1. (A)** Representative images showing the SOX2 (green) and ribosome biogenesis factor RSL24D1 (red) in control (WT) and mutant (V190G) day 15 organoids. Scale bar, 50 µm. **(B)** Quantification of the averaged nuclear RSL24D1 signal intensity of individual neuroepithelial bud at day 15 control (WT) and mutant (V190G) organoids. $*p < 0.05$, Unpaired t test with Welch's correction. n = 10 control and 9 mutant imaged regions from 2 independent batches. error bars are SEM.

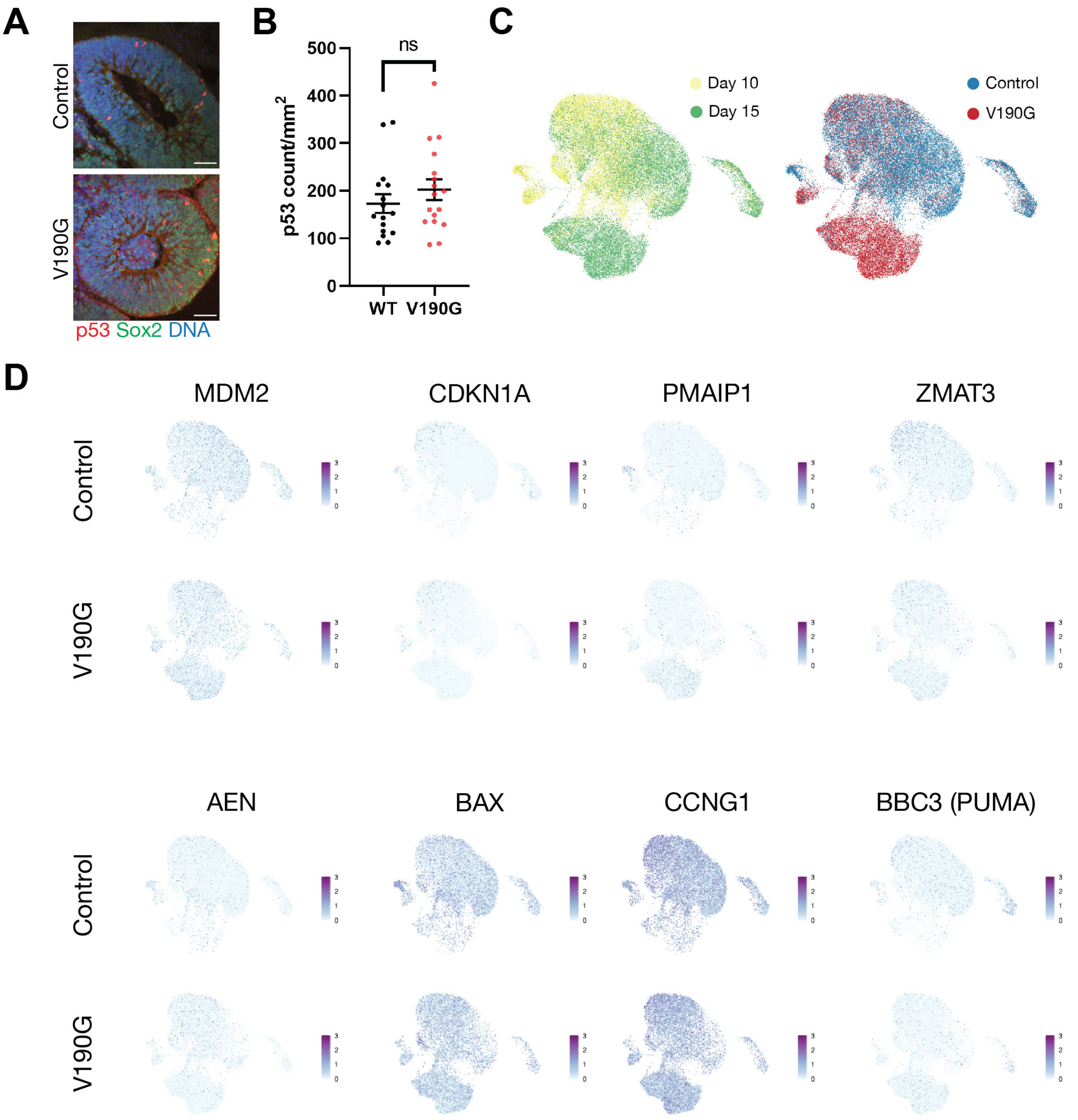


**Figure S5. Comparison of r-protein genes and p53 target gene expression in control and AIRIM organoids. (A)** Representative images showing the SOX2 (green) and p53 (red) in control (WT) and mutant (V190G) day 15 organoids. **(B)** Quantification of nuclear p53 signal of individual neuroepithelial bud at day 15 control (WT) and mutant (V190G) organoids. Unpaired t test with Welch's correction. n = 16 control and 17 mutant imaged regions from 3 independent batches. error bars are SEM. **(C)** UMAP embedding of a subset of the organoid development at days 10 and 15 colored by time point and genotype. **(D)** Expression of individual p53 target genes in cells from control (WT) and mutant (V190G) samples.

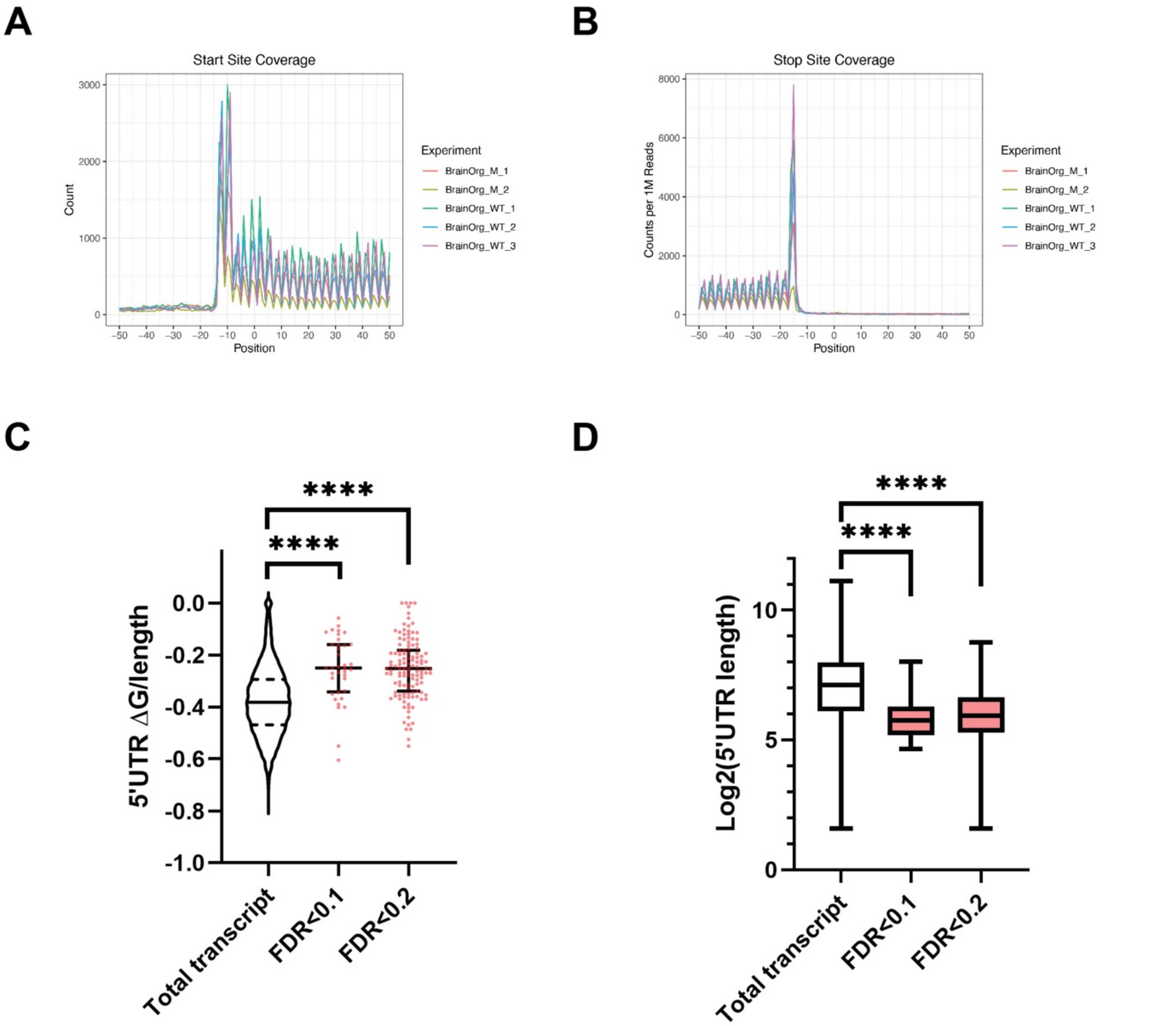


**E**

C1QBP
Gcuuccggcggcgccucaggucgcggggcgccu
aggccugggguuguccuuugcaucugcacguguu
cgcagucguuuccgcg

HSPD1
cccucacucgccgccgacgaccugucucgccgag
cgcacgccuugccgccgccccgcagaa

C12orf57
ccuuuccgcucccaggggcguugggaacgguug
uaggacguggcucuuuauucgugaguuuuccau
uuaccuccgcugaaccuagagcuucagacgcccu

**F**

| Motif | P-value | log P-pvalue | % of Targets | % of Background |
|---|---|---|---|---|
| 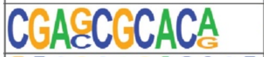 | 1e-4 | -9.935e+00 | 3.17% | 0.13% |
| 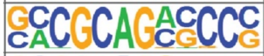 | 1e-3 | -7.949e+00 | 5.56% | 0.98% |
| 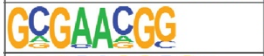 | 1e-3 | -7.405e+00 | 6.35% | 1.42% |
| 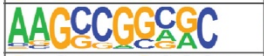 | 1e-2 | -6.080e+00 | 4.76% | 1.02% |
| 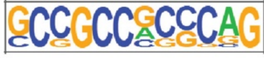 | 1e-2 | -5.510e+00 | 4.76% | 1.14% |

**Figure S6. Quality control and analysis of single-organoid ribo-seq analysis. (A)** The ribo-seq data exhibit high start site coverage across samples based upon meta-gene analysis of CDS regions. (**B**) The ribo-seq data exhibit high stop site coverage across samples based upon meta-gene analysis of CDS regions. (**C**) Violin/scatter dot plot quantifying TE changes relative to predicted 5'UTR free energy normalized to UTR length. ****$p < 0.0001$. Dunnett's T3 multiple comparisons test. n=7982(total), 37(FDR<0.1), 140(FDR<0.2) transcripts. (**D**) Box dot plot quantifying TE changes relative to log2 transformed 5'UTR length. ****$p < 0.0001$. Dunnett's T3 multiple comparisons test. n=7982(total), 37(FDR<0.1), 140(FDR<0.2) transcripts. (**E**) Example of TOP-like element containing transcripts identified in this study. (**F**) Motifs enriched in the 5'UTRs of *AIRIM*$^{V190G}$ sensitive transcripts are shown.

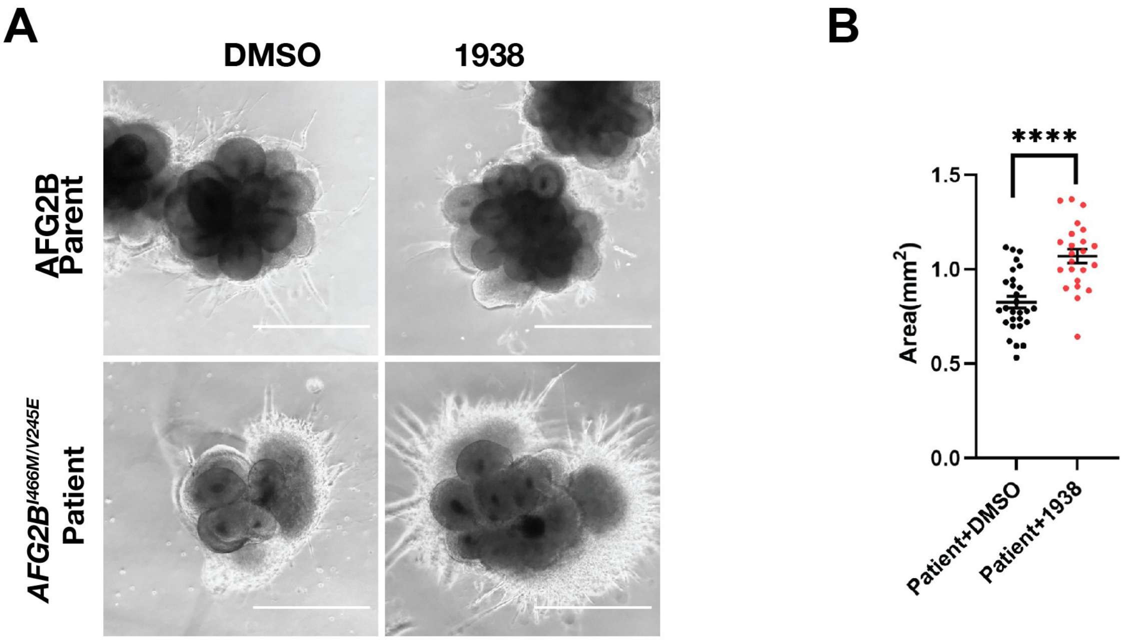


**Figure S7. Pharmacological activation of PI3Kα suppresses cerebral organoid growth defects caused by *AFG2B$^{I466M/V245E}$* genotype. (A)** Bright field images of day 15 control (parent) and patient (*AFG2B$^{I466M/V245E}$*) organoids treated with vehicle (DMSO) or 2µM PI3Kα activator UCL-TRO-1938 (1938). Scale bar, 1mm. **(B)** Quantification of size of day 15 patient organoids treated with vehicle (DMSO) or 2µM PI3Kα activator UCL-TRO-1938(1938). **** <0.0001. Unpaired t test with Welch's correction. n=28 DMSO and 23 UCL-TRO-1938 treated organoids, error bars are SEM. error bars are SEM.
