## Supplemental Table S1 for "An inappropriate decline in ribosome levels drives a diverse set of neurodevelopmental disorders"

### Extended Data Table 1. Summary of molecular and key clinical findings

| | Family 1 V:8, IV:21 | Family 2 II:3, II:4, II:5 | Family 3 II:1 | Family 4 II:2 | Family 5 II:1, II:3 | Family 6 II:1 | Family 7 II:3, II:5 |
|---|---|---|---|---|---|---|---|
| Consanguinity | Yes | No | Yes | Yes | Yes | Yes | Yes |
| Molecular genetics summary | | | | | | | |
| Chr1 g. position (GRCh37) | g.38152062T>G | g.38151999G>T | g.38155339G>A | g.38155339G>A | g.38155329C>G | g.38155329C>G | g.38148996A>C |
| Chr1 g. position (GRCh38) | g.37686390T>G | g.37686327G>T | g.37689667G>A | g.37689667G>A | g.37689657C>G | g.37689657C>G | g.37683324A>C |
| *AIRIM* c. position (NM_017850.3) | c.338A>C | c.401C>A | c.214C>T | c.214C>T | c.224G>C | c.224G>C | c.569T>G |
| AIRIM p. position (NP_060320.1) | p.(Y113S) | p.(S134Y) | p.(R72W) | p.(R72W) | p.(R75P) | p.(R75P) | p.(V190G) |
| Zygosity | Homozygous | Homozygous | Homozygous | Homozygous | Homozygous | Homozygous | Homozygous |
| In silico predictions | | | | | | | |
| phyloP [-19.0;10.9] | 3.55 | 7.60 | 4.54 | 4.54 | 0.50 | 0.50 | 5.46 |
| CADD | 26.4 | 28.0 | 32.0 | 32.0 | 24.7 | 24.7 | 25.2 |
| MT | D | D | B | B | B | B | B |
| PP2 | PrD | PrD | PrD | PrD | PrD | PrD | PrD |
| SIFT | D | D | D | D | T | T | D |
| REVEL | Uncertain | Damaging | Uncertain | Uncertain | Uncertain | Uncertain | Uncertain |
| Splicing Predictions | NA | NA | NA | NA | NA | NA | NA |
| ClinPred | Damaging | Damaging | Damaging | Damaging | Damaging | Damaging | Damaging |
| gnomAD v2.1.1 MAF* | 3.98e-6 | 0 | 7.08e-6 | 7.08e-6 | 0 | 0 | 0 |
| gnomAD v3.1.2 MAF* | 0 | 0 | 3.29e-5 | 3.29e-5 | 0 | 0 | 0 |
| All of Us Data Browser* | 0 | 0 | 8.00e-6 | 8.00e-6 | 0 | 0 | 0 |
| GME Variome* | 0 | 0 | 1.01e-3 | 1.01e-3 | 0 | 0 | 0 |
| Clinical summary | | | | | | | |
| Global developmental delay | +/+ | +/+/+ | NA | + | +/+ | + | +/+ |
| Intellectual disability | +/+ | +/+/NA | NA | + | +/+ | + | +/+ |
| Abnormal brain MRI | NA/NA | +/NA/NA | NA | + | NA/NA | + | NA/NA |
| Speech developmental delay | +/+ | +/+/+ | NA | + | +/+ | + | NA/NA |
| Motor developmental delay | +/+ | +/+/+ | NA | + | +/+ | + | NA/NA |
| Muscular hypotonia | +/+ | +/+/NA | NA | + | +/+ | + | NA/NA |
| Muscular hypotonia/limb spasticity/dystonia | +/+ | +/+/NA | NA | + | +/+ | + | NA/NA |
| Microcephaly | +/+ | +/+/NA | NA | + | +/+ | + | -/- |
| Seizures | -/+ | +/+/NA | NA | + | +/+ | + | -/- |
| Hearing impairment | NA/NA | +/+/NA | NA | NA | +/+ | - | +/+ |
| Vision impairment | NA/NA | +/+/NA | NA | + | +/+ | + | -/- |
| Feeding difficulties | +/+ | +/+/NA | NA | + | +/+ | + | NA/NA |
| Facial dysmorphism | +/- | -/-/NA | NA | + | -/- | + | -/- |
| Mortality | -/+ | -/-/- | + | - | -/- | - | -/- |

*no homozygous individuals were identified

Abbreviations: B, benign; D, Deleterious; DC, Disease causing; MAF, minor allele frequency; MT, MutationTaster; NA, Not available; PP2, PolyPhen-2; PrD, Probably damaging; T, Tolerated
